# Cell-Autonomous AR Dependence in Luminal Prostatic Epithelium Governs Survival and Lineage Plasticity

**DOI:** 10.64898/2026.02.13.705807

**Authors:** Dan Li, Naitao Wang, Wangxin Guo, Jude Owiredu, Woo Hyun Cho, Dana Schoeps, Shipeng Cheng, Hongjiong Zhang, Un In Chan, Chen Khuan Wong, Vaishnavi Reeya Callychurn, Hongsu Wang, Wenfei Kang, Ning Fan, H. Amalia Pasolli, Anurag Sharma, Anuradha Gopalan, Christopher E. Barbieri, Dong Gao, Ping Chi, Yu Chen

## Abstract

Prostate cancer resembles differentiated secretory luminal cells and shows cell-autonomous dependence on androgen receptor (AR) signaling, yet normal luminal cells are often considered dependent on paracrine stromal AR signaling. To resolve this, we conditionally deleted *Ar* in luminal acinar cells *in vivo*. *Ar*-deleted luminal cells persisted short-term, in contrast to the rapid regression observed after castration, but were impaired in regeneration and progressively lost. Their depletion was accompanied by replacement through basal-to-luminal differentiation of AR intact basal cells. Transcriptomic and chromatin profiling showed cell-autonomous suppression of the secretory program with induction of stemness, inflammatory, and epithelial-to-mesenchymal transition signatures after AR loss. Mechanistically, the MAP kinase pathway and downstream AP-1 transcription factors were activated and functionally validated, and MAP kinase inhibition selectively depleted AR-deleted luminal cells, indicating a compensatory survival pathway. These findings define intrinsic roles for luminal AR in maintaining differentiation, restraining plasticity, and sustaining regeneration and homeostatic turnover, providing a mechanistic basis for AR dependence in prostate cancer.

## Introduction

The prostate is a male reproductive secretory organ in which androgen receptor (AR) signaling is critical for proper development and function. AR is highly expressed in both epithelial and stromal cells. Seminal early studies showed that recombination of *Ar*-wildtype stroma with *Ar*-deficient epithelium generated prostate tissue capable of responding to androgens ^1^. Subsequent work demonstrated that epithelial-specific *Ar* knockout leads to increased proliferation, heightened inflammation, and impaired terminal differentiation, whereas stromal-specific *Ar* knockout results in decreased epithelial proliferation ^2–8^. Collectively, these findings support a model in which epithelial AR primarily regulates secretory function and terminal differentiation, while stromal AR governs epithelial lineage commitment and epithelial cell survival.

Recent single-cell RNA-seq studies in both human and mouse prostates consistently identify two major luminal epithelial subtypes^9, 10^. **Luminal acinar cells** localize to distal glandular regions and exhibit secretory activity. These cells have been variably labeled—for example, Luminal 1 ^11^, LumA/LumB ^12^, LumA/LumD/LumL/LumV ^13^, LumA/LumB/LumC ^14^, and Lum2/Lum4/Lum6–8 ^15^, – for clarity, we will refer to these as **Luminal 1**. **Luminal ductal and periurethral cells**, by contrast, are enriched in proximal ducts and as well as sparsely at the invagination tips of distal acini, show reduced secretory function, and display stem/progenitor-like signatures. These have been labeled as Luminal 2 ^11^, LumC ^12^, LumP ^13^, LumD ^14^, and Lum5 ^15^, we refer to these as **Luminal 2**. Androgen deprivation (such as castration) removes androgen ligand from stromal and epithelial compartments and reduces overall epithelial cell numbers and Luminal 1 cells preferentially. The remaining Luminal 1 cells shift transcriptionally with downregulating Luminal 1 genes and upregulating Luminal 2 markers ^11, 14, 15^. Notably, knockout of epithelial or luminal androgen receptor (AR) does not cause luminal cell loss, supporting a model in which androgen effects on luminal cells are mediated primarily through paracrine AR signaling from the mesenchyme rather than direct AR activity.

Prostate cancer is the most common malignancy in men, and prostate cancer cells typically display acinar histology and a transcriptomic profile resembling Luminal 1 epithelial cells of the benign prostate^16, 17^. In untreated prostate cancer, the androgen receptor (AR) represents a lineage-specific dependency, and androgen deprivation therapy (ADT)—which suppresses testicular testosterone production—has remained the cornerstone of treatment for advanced disease for more than 75 years^18^. The dependency of prostate cancer cells on AR signaling is cell intrinsic, and cancer cell–specific reactivation of AR signaling through AR mutation, amplification, alternative splicing, and diverse additional mechanisms commonly underlies the development of castration resistance ^19^. The seemingly conflicting roles of AR in normal luminal acinar cells versus prostate cancer cells have led to the hypothesis that tumorigenesis reprograms AR from regulating secretion and terminal differentiation to driving cell growth and survival^20–22^.

To understand the cell-intrinsic and paracrine roles of AR signaling in the normal prostate luminal cells, we compared castration with luminal cells-specific Ar knockout in the mouse prostate. Despite dramatic differences in histologic response, bulk and single-cell RNA-sequencing showed that castration and Ar knockout in Luminal 1 cells caused a remarkably similar transcriptional response, with cell-intrinsic downregulation of the secretory program and activation of stemness, inflammatory and epithelial-to-mesenchymal transition (EMT) programs.

Although Luminal 1 cells survived *Ar* loss in the short-term, they exhibited defects in regeneration following castration, developed endoplasmic reticulum stress accompanied by autophagy and mitophagy, and were ultimately replaced by basal cells over the longer term. Ar knockout led to activation of MAP kinase signaling and induction of downstream AP-1 transcription factors. Treatment with trametinib in vivo and expression of a dominant-negative AP-1 construct in 3D organoids impaired luminal cell survival and suppressed activation of the AR transcriptional signature, respectively.

## Results

### Cell intrinsic requirement of Ar for acinar differentiation in luminal cells

To study the cell autonomous effects of luminal-cell specific deletion of *Ar* in the prostate, we crossed ***T****mprss2-CreER^T2^*that is active in the prostate luminal cells^12, 23, 24^ with a *LSL-E**Y**FP* reporter^25^ (**TY**), with and without a conditional ***A****r* floxed allele^26^ (**TYA**). We administered tamoxifen to 8-week-old male mice and examined the prostates 2 weeks later (**Figure 1A**). To compare with Ar activity inhibition of all cell types in the prostate, we castrated a cohort of **TY** mice two weeks after tamoxifen and examined the prostates two weeks later (**TY-castrate**). We observed no significant change of prostate weight 2 weeks post-tamoxifen in **TYA** mice, whereas castration led to decreased prostate weight after two weeks (**Figure 1B**).

**Figure 1.**
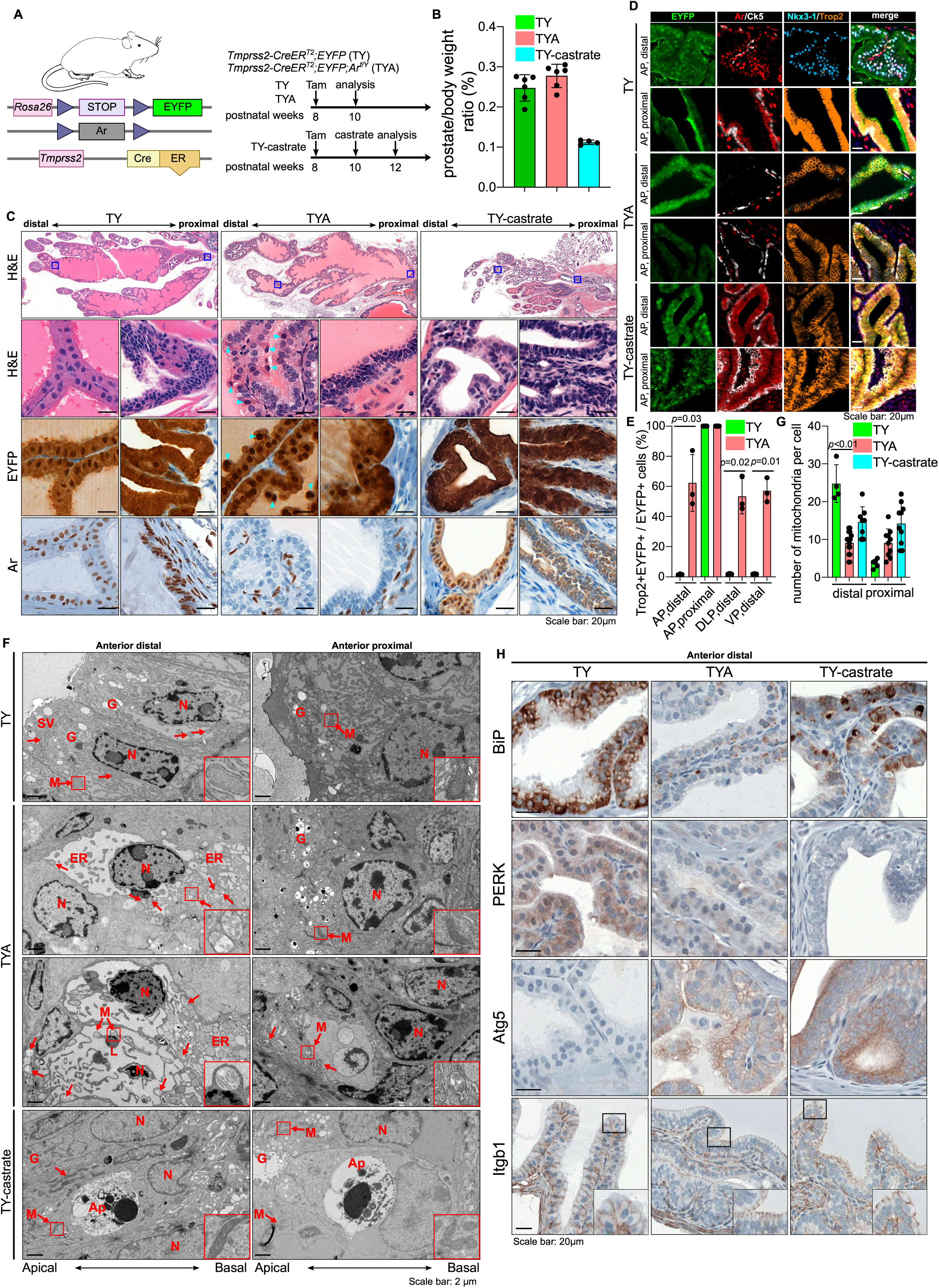
Histologic, immunohistologic and electron microscopic characterization of luminal Ar deletion and castration **(A)** Schematic of *Tmprss2-CreER^T2^; LSL-EYFP* control mice (TY) and *Tmprss2-CreER^T2^; LSL-EYFP; Ar^f/Y^* Ar knockout mice (TYA) and experiment strategies. Two doses of tamoxifen (3 mg ×2) were injected intraperitoneally with 48 hours interval. Castration surgery is performed to TY mice 2 weeks after tamoxifen injection (TY-castrate). **(B)** Mouse body and prostate weights 2 weeks after tamoxifen injection (TY and TYA), or 2 weeks after castration (TY-castrate). **(C)** Low magnification of hematoxylin and eosin (H&E) staining of anterior prostate lobe of TY, TYA and TY-castrate mice with boxes indicating magnified areas (top). High magnification of H&E and EYFP and Ar IHC of distal and proximal prostate (bottom). Arrowheads indicate necrotic nuclei from TYA mice. **(D)** Representative multiplex immunofluorescence (IF) of indicated proteins in distal and proximal regions of anterior prostates in TY, TYA and TY-castrate mice. **(E)** Quantification (percentage of Trop2^+^ EYFP^+^/EYFP^+^ cells) from distal and proximal anterior (AP), distal dorsal/lateral (DLP), and distal ventral (VP) prostate lobes from IF staining (mean ± s.d, two-tailed *t*-test). **(F)** Representative transmission electron microscopy (TEM) images of distal and proximal anterior lobes of TY and TYA mice 2 weeks after tamoxifen injection, and TY-castrate mice 2 weeks after castration. SV, Secretory vesicles; N, Nucleus; M, Mitochondrion; L, Lysosome; ER, Endoplasmic Reticulum; Ap, Apoptotic body; G, Golgi apparatus. **(G)** Quantification of number of mitochondria per luminal cell is quantified (mean ± s.d, two-tailed *t*-test). **(H)** Representative IHC staining of ER-associated proteins (BiP, PERK), autophagy-related gene (Atg5), and cell membrane protein Integrin beta 1 (Itgb1) in distal anterior prostates in TY, TYA and TY-castrate mice. Scale bar, 20 mm (panel C, D, G); 2 mm (panel F).

To quantify Ar deletion efficiency in luminal and basal cells of **TYA** prostates, we performed immunofluorescence against Ar, Ck8 and Ck5 in the distal region of anterior lobe (AP), lateral lobe (LP), dorsal lobe (DP) and ventral lobe (VP) and the proximal region of the anterior lobe. We observed highly efficient luminal Ar deletion except for the ventral lobe (**Figure S1A-B**). By immunohistochemistry, **TYA** prostates showed loss of Ar staining in luminal epithelial cells, but not basal or mesenchymal cells while **TA-castrate** prostates showed decreased nuclear Ar localization (**Figure 1C**, anterior lobe). Histologically, distal luminal acinar cells in **TYA** prostates exhibited nuclear dysplasia, characterized by enlarged, less basally localized nuclei with prominent nucleoli. There were numerous apically located pyknotic nuclei shedding into the lumen, indicative of increased cell death. A modest increase in proliferation, as assessed by Ki-67 staining, was also detected consistent with prior reports (**Figure 1C, S1C–D**)^3, 5^. In contrast, prostates examined two weeks after castration displayed marked reductions in weight, cellularity, and luminal secretions (**Figure 1C**). With both **TYA** and **TY-castrate**, while luminal acinar cells in the distal prostate exhibited histological changes in response to *Ar* knockout or castration, luminal ductal cells in the proximal prostate showed minimal histological alterations.

To assess changes in luminal cell differentiation induced by *Ar* knockout and by castration, we performed immunofluorescence for the Luminal 1 marker Nkx3-1 and the Luminal 2 marker Trop2 (*Tacstd2*). In intact mice, Trop2 labeled a small subset of luminal cells at the invagination tips of the distal prostate and all luminal cells in the proximal prostate. Nkx3-1 marked luminal cells only in the distal regions of all lobes except the ventral lobe and was absent in the proximal region (**Figure 1D–E, S1E**). Two weeks after *Ar* knockout or castration, distal luminal cells in all four lobes gained Trop2 expression and lost Nkx3-1 expression, adopting features characteristic of Luminal 2 cells. CD26 (*Dpp4*), a reliable marker of distal luminal cells localized to the apical surface, remained unchanged following *Ar* knockout (**Figure S1F**). We next used flow cytometry (FACS) to analyze cells from TY and TYA prostates. Epcam and CD49f distinguished luminal, basal, and stromal populations (**Figure S2A**). EYFP labeling was detected in most luminal cells (**TY**: 93.83 ± 2.08%; **TYA**: 88.93 ± 7.41%) (**Figure S2B**). Trop2 intensity formed a continuum and could not be cleanly divided into positive and negative populations. Using 95% of basal cells (which uniformly express Trop2) as a cutoff, the proportion of Trop2-positive luminal cells increased significantly in **TYA** compared to **TY** mice (**Figure S2A, C**). FACS-sorted Trop2-positive cells from **TY** and **TYA** prostates exhibited similarly higher organoid-forming efficiency compared to Trop2-negative cells indicating that Luminal 1 cells after Ar knockout exhibit similar organoid forming capacity as wild-type Luminal 2 cells (**Figure S2D–E**).

### Cell intrinsic requirement of Ar for mitochondrial and endoplasmic reticulum homeostasis

We observed nuclear dysplasia and increased cell death in distal luminal acinar cells after Ar knockout (**Figure 1C**). To examine the ultrastructure of luminal cells, we employed transmission electron microscopy (EM). Acinar luminal cells of the distal anterior lobe displayed well-developed, slightly dilated endoplasmic reticulum (ER) with electron-dense contents, consistent with active protein secretion, proximal luminal cells contain much less secretory vesicles and mitochondria (**Figure 1F**). In **TYA** mice, the ER of many cells exhibited loss electron-dense material and showed varying degrees of vacuolization, in some cases occupying nearly the entire cytoplasm. Ribosome attachment confirmed these vacuoles as ER-derived. Mitochondria were also affected in **TYA** luminal cells, exhibiting reduced matrix electron density and partial cristae loss (red arrows), compared with normal mitochondria in **TY** luminal cells. The mitochondria number per luminal cell also decreased in distal anterior prostate of TYA mice (**Figure 1G**). These cells appeared to represent progressive stages of ER vacuolization with plenty of secondary lysosomes, indicating mitophagy and autophagy. The ultrastructural features were not consistent with classical apoptosis, as nuclear fragmentation and apoptotic body formation were absent. In **TY-castrate** prostates, distal luminal cells exhibited decreased ER density, and some cells underwent classical apoptosis, featured with much slimmer and fewer mitochondria.

To further assess the ER density, we performed immunohistochemistry of ER resident chaperone BiP, the protein kinase R (PKR)-like endoplasmic reticulum kinase (PERK) (**Figure 1H, S1G**)^8, 27^. In TY mice, luminal acinar cells in distal anterior prostate exhibited strong staining consistent with their secretory function, while ductal luminal cells had much lower staining of both proteins. We observed decreased staining of BiP and PERK in both **TYA** and **TY-castrate** distal luminal epithelial cells, suggestive of compromised ER function. We performed Atg5 staining to mark autophagosomes^28^ and observed increased staining in luminal cells of **TYA** and **TY-castrate** mice, consistent with the increased autophagy in response to loss of Ar signaling. To assess cellular polarization, we performed staining against β1-integrin normally localized to basolateral membrane in polarized epithelia^29^. In **TYA** and **TY-castrate** mice, there was aberrant apical of β1-integrin together with more apically localized nuclei. Together, these findings indicate that *Ar* knockout drives a luminal cell-intrinsic shift toward a less differentiated, less secretory and more progenitor-like state.

### Acinar cell intrinsic Ar is required for androgen-mediated regeneration after castration

Androgen deprivation leads to rapid regression of the distal prostate, reaching a new equilibrium within four weeks, while androgen addback induces rapid regeneration^30, 31^. Previous studies using the *Nkx3-1-CreER^T2^* allele—which marks approximately 20% of luminal cells in the intact state and about 1% in the castrate state, referred to as castration-resistant Nkx3-1-expressing cells (CARNs)—suggested that *Ar* is generally dispensable for luminal regeneration but is important for the regenerative capacity of CARNs ^6, 7, 32^. Using the more efficient *Tmprss2-CreER^T^*^2^ allele, we investigated the role of luminal *Ar* in prostate regeneration following androgen addback. Two weeks after tamoxifen administration, mice underwent surgical castration, followed by implantation of dihydrotestosterone (DHT) pellets and EdU administration two weeks later (**Figure 2A**). Prostate weight was measured at 0, 3, 7, and 14 days after DHT addback. Compared to **TY** mice, **TYA** mice exhibited reduced prostate weight gain over time (**Figure 2B, S3A**). At two weeks post-DHT addback, **TY** prostates were fully regenerated and histologically indistinguishable from the pre-castration state, including restored Nkx3-1 expression. In contrast, **TYA** prostates showed impaired regeneration, resembling the regressed state and lacking Nkx3-1 expression (**Figure 2C**). Immunofluorescence analysis confirmed that in **TY** mice, Trop2 expression was lost, and Nkx3-1 expression was restored in the distal regions of all four lobes after regeneration. In contrast, **TYA** mice retained Trop2 expression and failed to re-express Nkx3-1 (**Figure S3B**). To assess proliferation in response to androgen addback, we analyzed EdU incorporation and found that **TYA** mice had fewer EdU-positive luminal cells at early time points (**Figure 2D–E**). Interestingly, in **TY** mice, EdU-positive cells were found throughout the prostate gland, whereas in **TYA** mice, EdU-positive cells were preferentially localized to glandular edges, suggesting a paracrine effect from stromal cells (**Figure 2D**). These data indicate that *Ar* plays a critical cell-autonomous role in regeneration.

**Figure 2.**
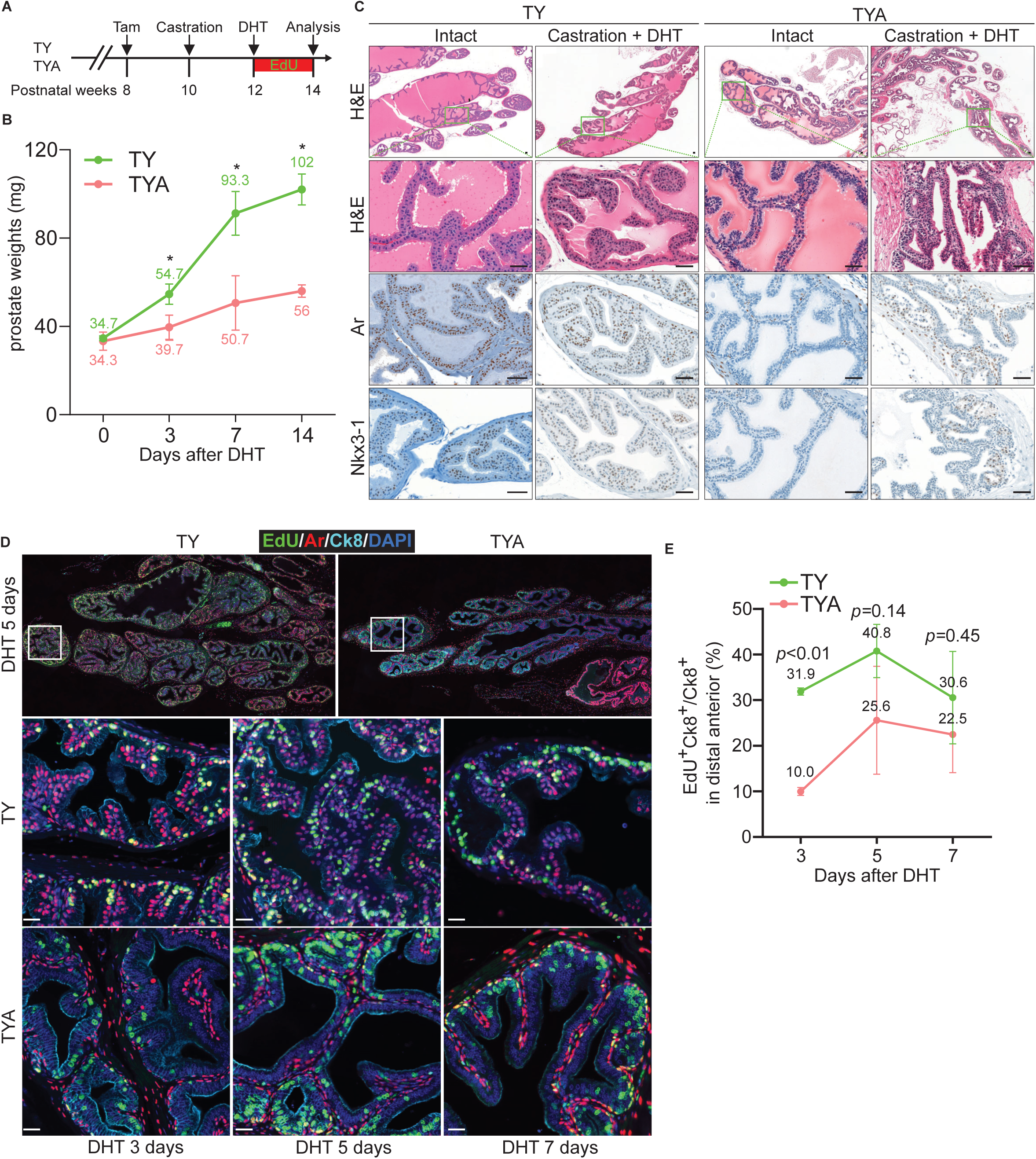
Ar loss attenuates luminal cell regeneration (**A**) Schematic showing timing of castration surgery, DHT pallet implantation and daily EdU administration. (**B**) Prostate weights are measured at 0, 3, 7 and 14 days after DHT transplantation. Asterisks indicate *p* value less than 0.05 (two-tailed *t*-test). (**C**) Representative H&E and Ar and Nkx3-1 IHC images of distal anterior prostates in TY and TYA mice after 2 weeks DHT transplantation. (**D**) Representative multiplex IF of EdU and indicated proteins are shown in distal anterior prostates in TY and TYA mice at 3, 5, 7 days after DHT transplantation. (**E**) Quantification of percentage of EdU^+^Ck8^+^/Ck8^+^ cells from IF staining. Data are presented as mean ± s.d. and analyzed with two-tailed *t*-test. Scale bar, 20 mm.

### Ar knockout luminal acinar cells cannot be sustained and are replaced by basal cell-derived luminal cells over time

Given the impaired regenerative capacity and evidence of cell stress on histology and electron microscopy of Ar-knockout luminal acinar cells, we investigated whether these cells exhibit reduced fitness and are progressively lost over time. Six months after tamoxifen administration, **TYA** prostates had regained a histologically normal appearance and maintained normal prostate weight (**Figure S4A-B**). Immunohistochemical analysis showed that distal luminal cells re-expressed Ar and lost EYFP expression. A more mosaic pattern was observed in the proximal prostate, characterized by a mix of **Ar** re-expression and **EYFP** loss. In contrast, luminal cells in **TY** prostates—whether intact or castrated—retained EYFP expression (**Figure S4B**). These findings suggest that luminal cells are stably maintained under both normal and castrate conditions; however, loss of Ar impairs their long-term fitness, resulting in their gradual replacement.

The restoration of Ar expression in luminal cells may stem from either the expansion of cells that escaped Cre-mediated Ar deletion or from a basal-to-luminal cell transition. To track both lineages in the prostate, we generated **TTR** mice by combining *Tmprss2-CreER^T2^*, *Trp63-DreER^T^*^2^, and the *Rosa26* Traffic Light Reporter (*R26-TLR*), which contains a *loxP*-flanked STOP cassette upstream of tdTomato and a *rox*-flanked STOP cassette upstream of ZsGreen^33, 34^. To evaluate the impact of Ar deletion, we introduced the *Ar*-floxed allele to create **TTRA** mice (**Figure 3A**). Two weeks after tamoxifen administration, immunofluorescence showed that most basal cells marked by Trp63 were labeled with ZsGreen, and most luminal cells marked by Ck8 were labeled with tdTomato in both **TTR** and **TTRA** prostates (**Figure 3B-C**). Using FACS quantification, we observed that ∼66% of luminal cells were tdTomato positive and ∼89% of basal cells were ZsGreen positive in both TTR and TTRA mice 2 weeks after tamoxifen administration (**Figure 3D, S5A)**.

**Figure 3.**
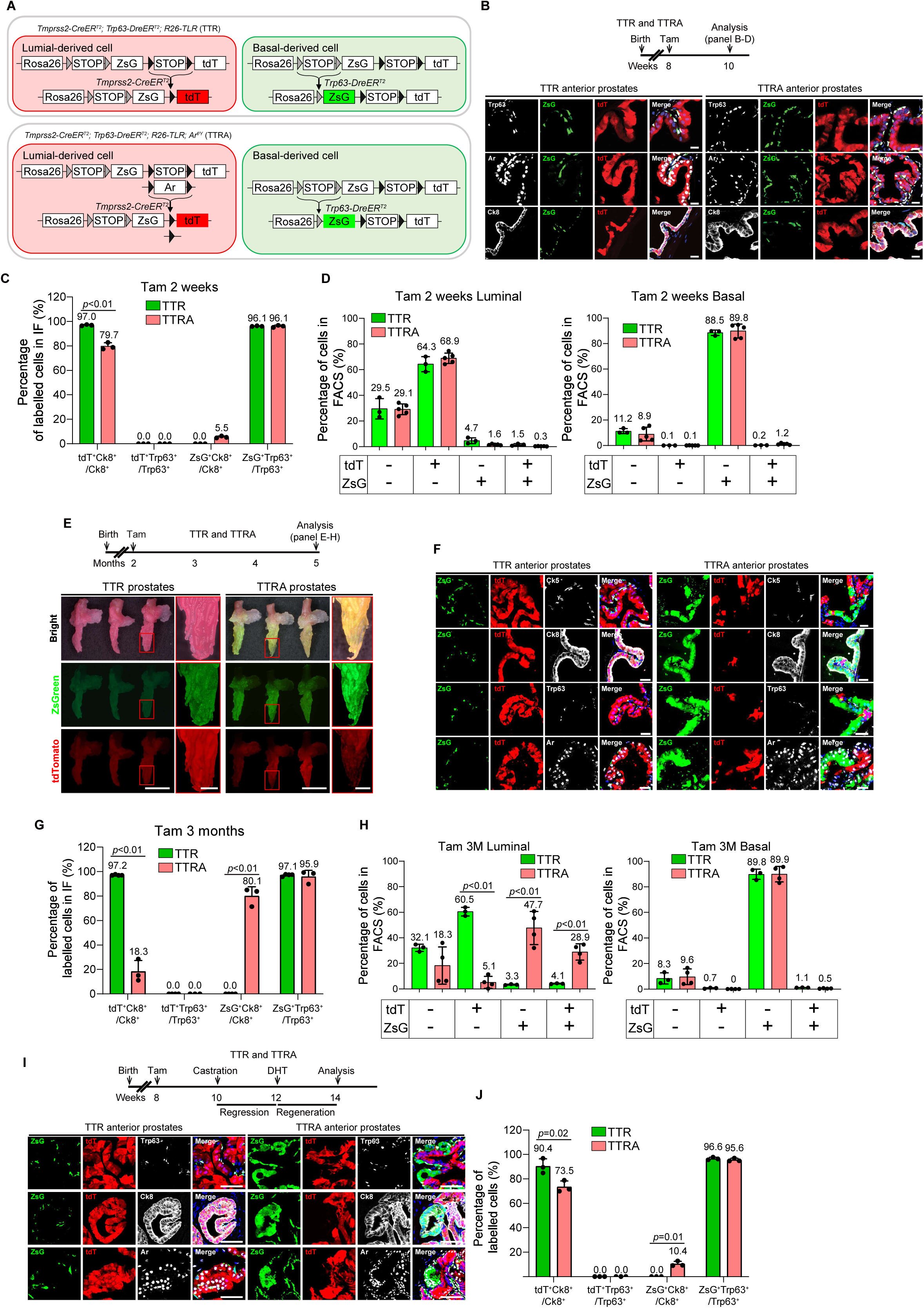
Ar loss induces luminal cell elimination and replacement with basal cell-derived Ar positive luminal cells (**A**) Schematic of lineage tracing *Tmprss2-CreER^T2^;Trp63-DreER^T2^*;*R26-TLR* (TTR) and *Tmprss2-CreER^T2^;Trp63-DreER^T2^*;*R26-TLR;Ar^f/Y^* (TTRA) mice to label *Tmprss2*-expressing cells by tdTomato (tdT) and label *Trp63*-expressing cells by ZsGreen (ZsG) after tamoxifen injection. (**B**) Representative multiplex immunofluorescence staining of indicated proteins are shown in distal anterior prostates of TTR and TTRA mice 2 weeks after tamoxifen injection. (**C**) Quantification (percentage of tdT^+^Ck8^+^/Ck8^+^, tdT^+^Trp63^+^/Trp63^+^, ZsG^+^Ck8^+^/Ck8^+^, ZsG^+^Trp63^+^/Trp63^+^ cells) from IF staining in panel B. Data is presented as mean ± s.d. (**D**) FACS based quantification of tdTomato and ZsGreen fluorescence in CD49f^Lo^, Epcam^Hi^ luminal cells and CD49f^Hi^, Epcam^Lo^ basal cells 3 months after tamoxifen administration. (**E**) Tissue fluorescent images show TTR and TTRA prostates 3 months after tamoxifen injection. (**F**) Representative multiplex IF staining of noted proteins in distal anterior prostates of TTR and TTRA mice 3 months after tamoxifen injection. (**G**) Quantification from IF staining in panel F. (**H**) Quantifications of each category of FACS data as in panel D. (**I**) Representative multiplex IF staining of noted proteins is shown in distal anterior prostates of TTR and TTRA mice 2 weeks after DHT transplantation. (**J**) Quantification from IF staining in panel I. Data is presented as mean ± s.d. (means are shown). Statistical tests are two-tailed *t*-test.

At three months post-tamoxifen, there was an overall decrease in tdTomato fluorescence and increase in zsGreen fluorescence of the whole prostate of **TTRA** mice (**Figure 3E**). Fluorescence microscopy showed that in **TTR** prostates, luminal cells continue to be labeled with tdTomato and basal cells with ZsGreen, suggesting that luminal and basal lineages are self-renewing (**Figure 3F**). In contrast, in **TTRA** prostates, among Ck8 labeled luminal cells the tdTomato-positive cells significantly decreased (TTR: 97.2%, TTRA: 18.3%) and ZsGreen-positive cells significantly increased (TTR: 0.0%, TTRA: 80.1%) (**Figure 3F-G**). Both ZsGreen-positive and tdTomato-positive luminal cells expressed Ar. These data suggest that three months after Ar-knockout in luminal cells, the luminal compartment in **TTRA** mice is mostly comprised of basal-derived luminal cells. FACS quantification revealed that ∼7% of luminal cells in TTR prostates were ZsGreen-positive whereas ∼77% of luminal cells in TTRA prostate were ZsGreen-positive (**Figure 3H**).

We crossed *Rosa26-CreER^T2^* with *Ar^flox^*(**R26A**) to delete Ar in all cells and compared with *Rosa26-CreER^T2^*(**R26C**) control mice (**Figure S4C**). At two weeks and 6 months after tamoxifen, **R26A** mice had decreased prostate weight similar to castrated mice (**Figure S4D**). Six months after tamoxifen administration, the prostate continued to exhibit a castrated histology. There were scattered stromal cells that stained positive for Ar while both basal and luminal cells remained Ar negative (**Figure S4E**). This data further supports that basal cells are the source of Ar-positive luminal cells after luminal-specific Ar deletion, and the basal to luminal lineage conversion is Ar dependent.

Given the defective regeneration of Ar knockout luminal cells, we investigated whether basal to luminal transition is stimulated by a cycle of castration-regeneration. In **TTR** prostates, ZsGreen expression remained confined to basal cells and tdTomato to luminal cells, indicating that this regenerative cycle did not promote lineage conversion. In contrast, **TTRA** prostates contained clusters of ZsGreen-positive luminal cells that have regained Ar expression, while most tdTomato positive luminal cells remained Ar-negative (**Figure 3I-J**). These findings suggest that, in the absence of luminal Ar, basal-to-luminal differentiation is enhanced during regeneration, likely compensating for the compromised luminal population. Overall, our data indicate that Ar is essential for the long-term maintenance and regenerative capacity of luminal epithelial cells.

### Cell-intrinsic Ar mediates acinar transcriptional program and inhibits inflammatory signaling in luminal cells

To identify cell-intrinsic and paracrine effects of Ar signaling on prostate luminal epithelial cells, we performed RNA-seq on FACS-sorted EYFP-positive, Epcam^+^CD24^high^ luminal cells from **TY**, **TY-castrate** (5 days after castration), and **TYA** mice two weeks after tamoxifen administration (**Figure S6A**). Principal component analysis (PCA) revealed the triplicates were clustered together. The first principal component (PC1), accounting for over 50% of the variance, showed similar directional changes in TYA and TY-castrate (**Figure S6B**). At the gene level, castration resulted in a balanced number of significantly upregulated and downregulated genes (fold change > 2, FDR < 0.05), whereas Ar knockout led to a larger number of significantly upregulated genes (**Figure 4A**). Among genes significantly altered by both perturbations, expression changes were highly correlated (**Figure 4B**). Genes significantly perturbed by castration only were mostly downregulated by castration whereas genes significantly perturbed by Ar knockout only were upregulated. We conducted Gene Set Enrichment Analysis (GSEA) comparing **TY-castrate** vs. **TY** and **TYA** vs. **TY** using Hallmark and Curated gene sets from the Molecular Signatures Database, along with custom gene sets for prostate Luminal Distal and Luminal Proximal cells^10^. The LUMINAL_DISTAL gene set, comprised of genes defining in Luminal 1 acinar cells was the most downregulated in both castrated and Ar knockout samples (**Figure 4C, S6C, Table S1**). Several metabolic gene sets—including those related to amino acids, lipids, and mitochondrial functions were also downregulated under both conditions. Conversely, gene sets upregulated in both castration and Ar knockout include the LUMINAL_PROXIMAL gene set comprised of genes defining Luminal 2 ductal cells, inflammatory pathways (e.g., interferon and TNF signaling), and epithelial-to-mesenchymal transition (EMT) signatures (**Figure 4C, S6C, Table S1**). Cell cycle and extracellular matrix gene sets were downregulated in castration but upregulated in Ar knockout samples, aligning with the rapid prostate regression observed post-castration but increased Ki-67 after Ar knockout. We examined representative genes from these sets, including Luminal Distal markers (*Nkx3-1, Pbsn, Tgm4*), the Luminal Proximal markers (*Clu*, *Tacstd2*), inflammatory genes (*Fosl1, Foxl2, Junb, Atf3, Nr4a1, Egr1, Irf9*, *Rela*), Yap/Taz downstream genes (*Ccn1* and *Ccn2*) and cell cycle regulators (*Mki67, Cenpf, Cdc20, Ube2c*), which illustrate these transcriptomic shifts (**Figure 4D**).

**Figure 4.**
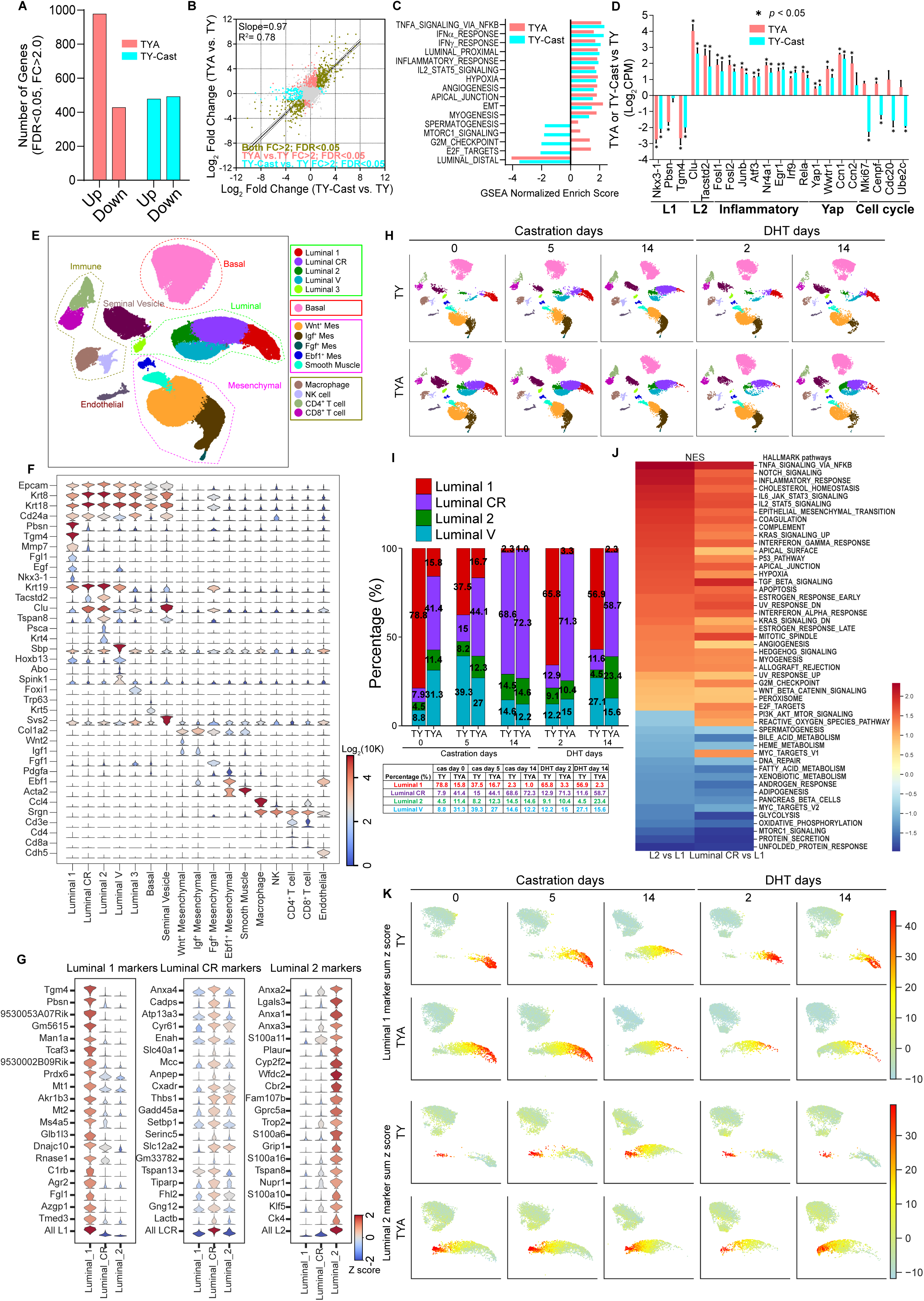
Transcriptomic survey shows Ar intrinsically regulating luminal cell functions and regeneration (**A**) The number of significant up-regulated and down-regulated genes from RNA-seq of sorted luminal cells of TYA and TY-castrate compared with TY prostates. (**B**) Scatter plot and correlation analysis of differentially expressed genes of TYA compared with TY, TY-castrate compared with TY by bulk RNA-seq using sorted luminal cells. Fit line is generated from comparing genes changed by both castration and Ar-knockout. (**C**) GSEA normalized enrichment score of representative gene sets comparing TYA with TY and TY-castrate with TY. (**D**) The expression changes of noted luminal distal, proximal and cell cycle related genes base on RNA-seq. (**E**) Uniform manifold approximation and projection (UMAP) of single-cell RNA-seq (scRNA-seq) profiles colored by Leiden clustering of subsets. (**F**) Violin plots of cell markers by cluster. The color in the violin plots indicates the median normalized expression level of genes. (**G**) Violin plots of Luminal 1, Luminal 2 and Luminal CR cell markers in each cell cluster. (**H**) Separate UMAP of triplicate samples by mouse and treatment (**I**) Bar graph (top) and percentage (bottom) of quantification of the four luminal cell subsets in the indicated mice and treatment. (**J**) Heatmap of GSEA normalized enrichment score of the Hallmark genes sets in Luminal 2 and Luminal CR compared with Luminal 1. (**K**) Summed z score of Luminal 1 and Luminal 2 signature genes separately for each sample on UMAP.

To characterize cell intrinsic and paracrine Ar singling in luminal subpopulations, we performed single-cell RNA sequencing (scRNA-seq) using freshly dissociated whole prostates of intact, castrated, and DHT treated **TY** and **TYA** mice, each condition contained 3 biological replicates labeled with hashing tags. We profiled 92,371 individual cells, which can be clustered to 17 distinct clusters (**Figure 4E**). The luminal and basal populations could be distinguished by luminal markers *Ck8, Ck18, Cd24a* and basal markers *Ck5, and Trp63* respectively (**Figure 4F**). We identified 5 distinct luminal epithelial clusters were defined with published marker genes: Luminal 1 (*Pbsn*^high^, *Tgm4*^high^, *Mmp7*^high^, *Fgl1*^high^ and *Nkx3-1*^high^); Luminal 2 (*Tacstd2*^high^, *Clu*^high^, *Psca*^high^, and *Ck4*^high^); Luminal 3 (*Foxi1*^high^); Luminal V (*Sbp*^high^, *Hoxb13*^high^, *Abo*^high^, and *Spink1*^high^, Ventral prostate) and **c**astration-**r**esistant luminal cells Luminal CR (**Figure 4F**). Differential expressed genes between Luminal 1, Luminal CR and Luminal 2 showed that Luminal CR had Luminal 1 and Luminal 2 mixture gene expression pattern (**Figure 4G**).

In intact **TY** mice, Luminal 1 cells comprised the majority (∼79%) of luminal cells. After castration, there was progressive loss of Luminal 1 cells to just ∼2% and gain of Luminal CR cells to 69% by two weeks. DHT addback restored Luminal 1 cell population as early as two days (**Figure 4H-I**). In intact **TYA** mice two weeks after tamoxifen administration, Luminal CR was the dominant luminal cluster in intact state and maintained during castration and DHT addback. Luminal 1 cells decreased after castration and only very few remained after DHT treatment (**Figure 4H-I**). There was broad overlap in Luminal CR cluster in intact **TYA** mice and castrated **TY** mice, indicating that the transcriptional changes induced by castration in Luminal 1 cells are largely cell intrinsic.

We performed GSEA using HALLMARK gene sets comparing Luminal 2 and Luminal 1 only in intact **TY** mice (**Figure S6D**), or comparing Luminal CR and Luminal 2 with Luminal 1 in all intact, castrated and DHT treated **TY** and **TYA** samples (**Figure 4J**). Luminal CR and Luminal 2 shared upregulated gene sets associated with inflammatory signaling (i.e., TNF-NFkB and interferon-JAK-STAT) pathways, RAS signaling pathways and epithelial mesenchymal transition (EMT) pathways. Luminal CR and Luminal 2 shared downregulated gene sets were metabolism and protein secretion. There was high concordance between gene expression changes observed after Ar knockout in bulk and single cell RNA-seq analysis (**Figure 4C, 4J, S6C, Table S2**).

We calculated summed z score of Luminal 1 and Luminal 2 signature genes and plotted the scores on UMAP consisting of all luminal cells. Luminal CR had intermediate levels of Luminal 1 score and Luminal 2 score (**Figure 4K**). The Luminal 1 score decreased after castration and restored after DHT treatment, and the Luminal 2 score had an opposite transition in TY mice, but Luminal 1 score decreased after castration and could not restore after DHT treatment in TYA mice (**Figure 4K**).

In the stromal compartment, there were 5 mesenchymal clusters can be identified (*Wnt*^+^, *Igf*^+^, *Fgf*^+^, *Ebf*^+^, and smooth muscle). The *Wnt*^+^-Mesenchymal decreased after castration and restored after DHT addback, the *Igf*^+^-Mesenchymal increased after castration and reduced after DHT addback (**Figure S7A**). The marker genes in each cluster (*Wnt2, Igf1, Fgf1, Pdgfa,* and *Rspo3*) and the extracellular matrix (ECM) genes (*Col1a1*, *Col1a2, Col4a1, Col4a2, Col6a1, Col6a2*) all decreased after castration and restored after DHT addback (**Figure S7B-C**). Ar knockout in luminal cells did not induce significant transcriptional changes in basal or mesenchymal cells (**Figure S7D**). We independently replicated this experiment using anterior prostate only and observed Luminal 1 to Luminal CR shift with both castration and with Ar knockout (**Figure S8A-C**). The Luminal V population disappeared when only the anterior prostate was used, confirming that these luminal cells are from the ventral prostate. These data indicate that while luminal Ar knockout does not lead to prostate regression or expression changes in stromal and basal cells, luminal cells exhibit similar expression changes as castration suggesting that these changes are largely cell intrinsic and independent on stromal growth factor mediated prostate regression.

### Activation of AP-1 transcription factors can partially account for upregulated pathways in response to Ar loss

To identify putative transcriptional mediators after Ar inhibition, we examined changes in the chromatin accessibility landscape using Transposase-Accessible Chromatin sequencing (ATAC-seq) on FACS-sorted EYFP-positive, Epcam^+^CD24^high^ luminal cells from **TY**, **TYA**, and **TY-castrate** (5 days after castration) mice two weeks after tamoxifen administration (**Figure S6A**). We identified 8825, 9480, and 10059 reproducible promoter peaks and 19645, 34272, and 31778 reproducible enhancer peaks in **TY**, **TYA**, and **TY-castrate** respectively. The promoter peaks were largely overlapping while the enhancer peaks exhibited more changes between conditions, leading us to focus on enhancers for downstream analysis (**Figure S9A**).

We observed an overall concordance in change of ATAC signal by Ar knockout and by castration, with fewer Ar-knockout and castration specific changes (**Figure S9B**). In peaks that were commonly lost after Ar knockout or castration, we identified ARE and the composite FOXA1:AR as the most enriched motifs (**Figure 5A-B, Table S3**). In peaks that were commonly gained after Ar knockout or castration, we identified AP-1 and FOXA1 as the most enriched motifs (**Table S4**). Comparing Ar knockout with castration, we found the FOXA1:AR motif to be most enriched among Ar knockout decreased peaks and the NFκB motif as enriched among Ar knockout increased peaks (**Figure 5B, Table S5-6**). We next performed digital genomic footprinting analysis of ATAC-seq data to define transcript factor binding sites within peaks^35^. We observed highly concordant changes induced by Ar knockout and by castration, with increased binding of AP-1, Maf, and Nfe2 family transcription factors that are known to mediate stress and inflammatory responses (**Figure 5C, Table S7**). NFκB motif was specifically enriched in **TYA** and not **TY-castrate**.

**Figure 5.**
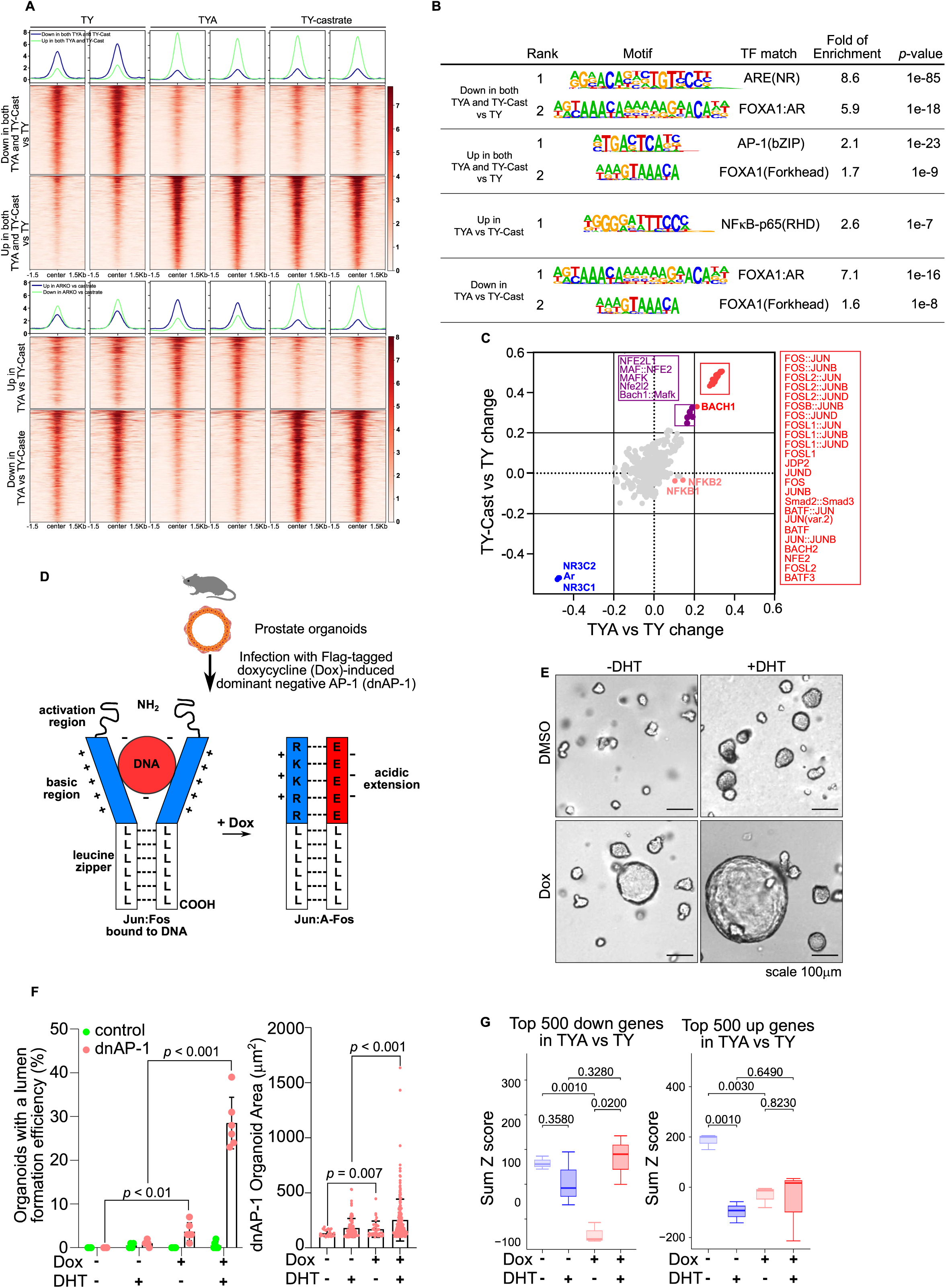
ATAC sequencing shows that Ar loss or castration increases AP-1 activity and AP-1 inhibition promotes luminal cell organoid formation efficiency and Ar target genes expression (**A**) ATAC-seq heatmap and profile of significantly up and downregulated enhancer peaks in TYA and TY-castrate compared with TY (top), and the up-regulated and down-regulated enhancer peaks in TYA compared with TY-castrate. (**B**) The enriched motifs categories of peaks in panel A are shown. (**C**) Correlation of differential binding scores based on TOBIAS ATAC footprinting analysis of TYA compared with TY and TY-castrate compared with TY. (**D**) Schematic of doxycycline inducible dnAP-1 expression in prostate cells. (**E**) Six-day organoid formation assay modulating AP-1 inhibition and Ar activation with 1mg/ml Dox and 10nM DHT addition respectively. (**F**) Quantification of percent of organoids with a lumen and organoid area from panel E. Data are presented as mean ± s.d. and analyzed with two-tailed *t*-test. (**G**) The transcriptional profiling of dnAP-1 expressed organoid cells is analyzed by RNA-seq. The sum Z-scores of top 500 down-regulated or up-regulated genes in TYA compared with TY in dnAP-1 expressed organoid cells are shown.

The dramatic gain in chromatin accessibility at AP-1 binding sites following *Ar* knockout and castration suggests that AP-1 family members may regulate genes that are induced upon Ar inhibition. In prostate organoids, we previously showed that DHT treatment decreases accessibility at AP-1 motifs, indicating a cell-intrinsic antagonism between Ar and AP-1 activity that can be modeled in vitro^21^. AP-1 transcription factors are formed by leucine-zipper dimers of Jun, Fos, and Atf family transcription factors. The complexity, diversity, and compensation of AP-1 family proteins make clean genetic depletion studies difficult. To globally inhibit AP-1 function, we introduced a doxycycline-inducible dnAP-1, a dominant negative AP-1 protein consisting of an acidic amphipathic protein sequence appended onto the N-terminus of the FOS leucine zipper, replacing the normal basic region critical for DNA binding. dnAP-1 dimerizes with endogenous AP-1 transcription factors, abrogating AP-1 activity across multiple family members (**Figure 5D**)^36, 37^. Expression of dnAP-1 in genetically normal mouse prostate organoids was confirmed by immunoblotting using a FLAG antibody (**Figure S9C**). The expression of Jun family genes, Jun, Junb and Jund did not change, but the Fos family genes Fosl1, Fosl2, Fosb decreased after dnAP-1 expression (**Figure S9C**).

AP-1 inhibition increased luminal features in organoids, including the number and size of organoids with lumen formation and Ck8 positive luminal cells (**Figure 5E-F, S9D**). We next examined the effect of Ar and AP-1 modulation in organoids on the genes affected by Ar knockout in prostate luminal cells *in vivo*. Among genes downregulated after Ar knockout (i.e., Ar-target genes), we observed that AP-1 inhibition led to increased DHT-mediated regulation. Genes upregulated by Ar knockout, were suppressed by DHT in organoids recapitulating the Ar suppressive effect. Notably, when dnAP-1 was expressed, these genes were inhibited in DHT-free conditions, suggesting that AP-1 transcription factors mediate transcriptional upregulation after Ar inhibition. Together, these studies show that AP-1 factors act to oppose an AR-driven luminal acinar transcriptional program in prostate epithelium.

### Activated MAP kinase signaling is a survival pathway after Ar knockout in luminal cells

We observed that AP-1 and NFκB transcription factors are transcriptionally upregulated after Ar knockout and castration (**Figure 4D**). Beyond expression, the activity of these factors is governed by signaling cascades. We thus examined activation of the MAP kinase pathway known to phosphorylate multiple AP-1 factors, phosphorylation of downstream AP-1 factors Atf2 and Jun, phosphorylation of Stat3 downstream of the JAK-STAT signaling pathway and localization of the NF-κB subunit p65 using IHC.

In the distal prostate at baseline, we observed occasional Erk phosphorylation in the basal cells and Atf2 phosphorylation in the stromal and basal cells and no evidence of Jun or Stat3 phosphorylation. In the proximal prostate, Atf2 phosphorylation was detected in the luminal cells without observable changes in other markers. After luminal Ar knockout, we observed robust increase in Erk phosphorylation and downstream Jun phosphorylation especially in the distal luminal cells. There was also appreciable increase in Stat3 phosphorylation and p65 protein levels in both distal and proximal luminal cells (**Figure 6A-B**). These data suggest that Atf2 may mediate baseline AP-1 activity in proximal L2 luminal cells but AR knockout and castration further activates multiple signaling pathways in a cell intrinsic manner.

**Figure 6.**
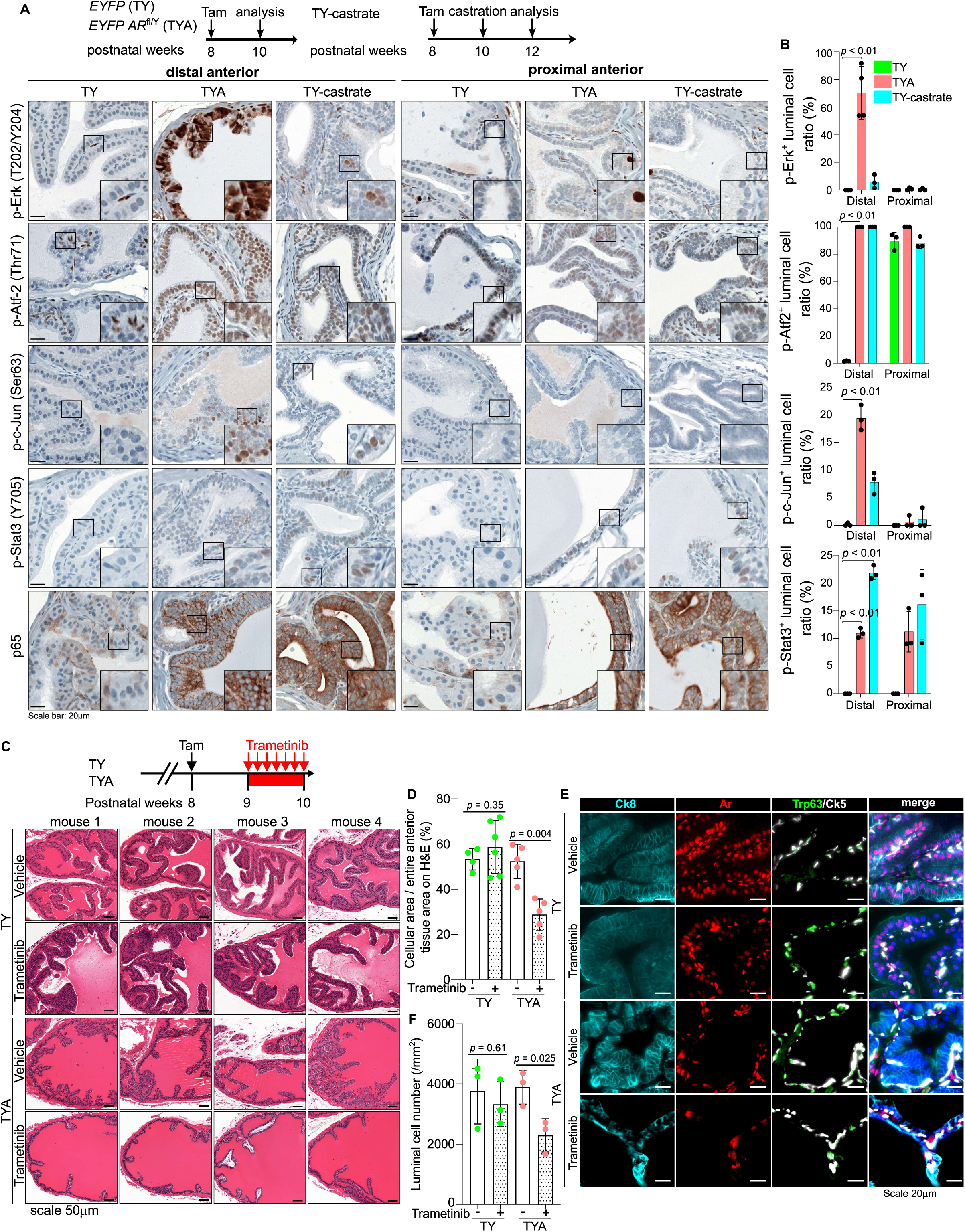
Ar loss induces AP-1/Erk activation, and trametinib decreases luminal cell survival *in vivo*. (**A**) Representative IHC staining of indicated proteins in distal and proximal anterior prostates in TY, TYA and TY-castrate mice. (**B**) Quantification from IHC staining in panel A. Positive cell numbers were quantified manual by eye and averaged on 3 representative areas on IHC images from distal or proximal anterior prostate of each mouse. At least three mice were used for each condition. Data is presented as mean ± s.d. (two-tailed *t*-test). (**C**) One week after Tamoxifen injection, the TY and TYA mice are treated with vehicle or trametinib (1 mg/kg daily by oral gavage) for 1 week before sacrifice. Representative H&E staining shows the histological changes of distal anterior prostates in TY and TYA mice after Trametinib treatment. (**D**) Quantification of the cellular area percentage of the entire anterior tissue area on H&E-stained slides in panel C using Fiji ImageJ software. Data is presented as mean ± s.d. (two-tailed *t*-test). (**E**) Representative IF staining of noted proteins in distal anterior prostates in TY and TYA mice after Trametinib treatment. (**F**) Quantification of the Ck8^+^ luminal cells on IF-stained slides in panel E. Data is presented as mean ± s.d. (two-tailed *t*-test).

Given the dramatic activation of Erk phosphorylation in response to Ar knockout, we determined the response to treatment with the Mek inhibitor trametinib. We treated TY and TYA mice with 1 mg/kg trametinib daily for one week starting from one week after tamoxifen injection. Although the body weights did not change significantly in each group, the prostate weights of TYA mice significantly reduced after trametinib treatment (**Figure S10A**). Immunohistochemistry showed that trametinib treatment inhibited Erk and Jun phosphorylation but did not affect Atf2 phosphorylation in TYA distal prostates (**Figure 6C**). Trop2 expression still maintained in distal anterior prostate of TYA mice after trametinib treatment (**Figure S10B**).

Histologically, trametinib led to loss of distal prostate glandular structure and infoldings readily apparent on H&E stains (**Figure 6D**). Quantification Ck8^+^ luminal cells showed that trametinib treatment decreased the number of distal luminal cells specifically in TYA mice and Ck8 immunofluorescence highlight a decrease in the cytoplasmic volume of luminal cells (**Figure 6D-E**). The remaining luminal cells were flattened with little cytosol, best appreciated on Ck8 IF staining. These data suggest that Ar knockout cells are dependent on MAP kinase signaling for survival.

## Discussion

In the prostate, AR is abundantly expressed in both stromal and epithelial cells and androgen signaling governs development, growth, differentiation and secretion. Cycles of castration and androgen addback leads to regression and regeneration of the prostate, where secretory luminal epithelial “Luminal 1” cells of the distal acini are most affected^38, 39^. Lineage tracing studies have shown that the luminal and basal compartments are largely self-sustained over time and through cycles of castration and regeneration, although inflammation and oncogene expression can lead to basal-to-luminal differentiation^32, 34, 40, 41^. Approximately two weeks after castration, the mouse prostate reaches a new equilibrium of the regressed state where the remaining luminal epithelial cells adapt a gene expression program similar to “Luminal 2” cells found in the proximal periurethral prostate^10–12^. These more stem-like, castration-resistant cells are capable of rapid proliferation followed by differentiation upon androgen addback. The epithelial regression and regeneration response is due to stromal AR signaling, as recombination of *Ar*-wildtype stroma with *Ar*-deficient epithelium generated prostate tissue capable of responding to androgens^1^ and epithelial-specific *Ar* knockout did not lead to regression^2, 3^.

The apparent Ar-independence of distal luminal epithelial cells seems to contrast with human prostate cancer which exhibit acinar adenocarcinoma histology, express secretory components and the transcription factor NKX3.1, similar to benign Luminal 1 cells. Clinical and experimental observations unequivocally demonstrate that the majority of prostate cancers exhibit cell-autonomous dependence on AR. Prostate cancer cell lines and xenografts model AR dependence and clinical castration resistance is frequently driven by tumor-intrinsic reactivation of AR through gene amplification, ligand-binding domain mutations, and splice variants^19^.

To address this discrepancy, we compared castration with a luminal epithelial-specific *Ar* knockout. Consistent with previous studies, castration led to rapid prostate regression that reached a new equilibrium within two weeks. In contrast, *Ar*-knockout luminal epithelial cells did not undergo rapid regression; instead, they exhibited enlarged nuclei and slightly increased proliferation^3, 5^. Despite these histological differences, the regressed luminal cells after castration and the *Ar*-knockout luminal cells shared a largely overlapping transcriptional and chromatin landscape changes: 1) Downregulation of secretory identity: There was a loss of secretory products and the protein secretion machinery itself, characterized by decreased endoplasmic reticulum (ER) density and the loss of AR binding sites. 2) Upregulation of Stress Signatures: We observed an increase in inflammatory and EMT (epithelial-mesenchymal transition) signature genes, accompanied by a gain of AP-1 binding sites. This state approximates the stem-like “Luminal 2” cells typically residing in the proximal periurethral zone. These findings indicate that the stereotypic transcriptional and epigenetic responses shared by castration and *Ar*-knockout are direct, cell-autonomous responses to the loss of AR signaling, rather than a selective loss of the acinar “Luminal 1” cell type.

We also identified critical differences between castration and *Ar* knockout. *Ar* knockout, especially in acinar Luminal 1 cells, caused profound defects in regeneration upon androgen add-back. These cells were gradually lost, replaced by basal cells, and exhibited markers of mitophagy and autophagy. These results indicate that AR is cell-autonomously required for luminal acinar cell survival, but on a longer timescale than the rapid regression seen with systemic castration. Prior studies suggested that *Ar*-deleted luminal cells can regenerate^6, 7^. While we observed an early proliferative burst in *Ar*-knockout cells after androgen add-back, consistent with others, this did not fully restore the epithelium. We speculate that more complete luminal *Ar* deletion using the *Tmprss2-CreERT2* allele, combined with longer follow-up, revealed this survival defect.

Both *Ar* knockout and castration activated multiple signaling pathways—STAT, NFκB, and MAP kinase (MAPK)—with increased chromatin accessibility at AP-1 and NFκB sites. Notably, MAPK activation was more pronounced in the *Ar*-knockout model. While normal Luminal 2 cells also exhibit AP-1 accessibility and phosphorylated Atf-2, they lack phosphorylated Erk, c-Jun, or Stat3. This prompted us to investigate MAPK and downstream AP-1 activity as potential survival mechanisms for luminal cells following AR loss. In organoids, AP-1 signaling was essential for inducing *Ar*-knockout–responsive genes. In vivo, MEK inhibition led to selective loss of luminal cells after *Ar* knockout. These findings highlight an interplay between AR and AP-1–driven transcription during prostate epithelial lineage specification.

Our work has implications for understanding human prostate cancers. Neoadjuvant androgen deprivation therapy (ADT) trials show minimal clinical response after two weeks, with responses continuing to evolve over six months—a timeline distinct from the rapid two-week involution seen in mice^42–44^. Our data suggests that the cell-autonomous AR dependence observed in prostate cancer is fundamentally rooted in the biology of normal luminal acinar cells, which are lost over a longer time period after *Ar* knockout, potentially paralleling clinical tumor responses.

Castration-resistant prostate cancer (CRPC) often emerges following prolonged ADT and can reactivate AR signaling or acquire lineage plasticity in which AR signaling becomes dispensable. We recently identified a lineage-plastic CRPC subtype, CRPC-SCL, characterized by inflammatory signaling and driven by AP-1 and YAP/TAZ pathways^45^. These same pathways arise early during lineage plasticity in mouse prostate cancer driven by *Trp53* and *Rb1* loss. Our findings suggest that normal acinar luminal cells are programmed to activate these pathways after AR inhibition^46^. These pathways support short-term survival but are insufficient for long-term maintenance, and loss of tumor suppressors such as p53 and RB can convert this transient state into one capable of lineage plasticity and renewed growth.

## Methods and Materials

### Mouse studies

Mouse experiments except those depicted in **Figures 3, S5, and S8** were conducted under protocol 11-12-027 approved by Institutional Animal Care and Use Committee (IACUC) of Memorial Sloan Kettering, New York. Mouse experiments shown in **Figures 3, S5, and S8** were administered under the guidelines of the Animal Care and Use Committee at the Center for Excellence in Molecular Cell Science, Chinese Academy of Sciences (CAS).

*Tmprss2-CreER^T2^-IRES-nlsEGFP* (*Tmprss2^tm1.1(cre/ERT2)Ychen^*, MGI:5911389)^23^, *LSL-EYFP* (B6.Cg-*Gt(ROSA)26Sor^tm3(CAG-EYFP)Hze^*/J, MGI:J:155793)^25^, *Trp63-DreER^T^*^247^, and *Rosa26-TLR*^33^ mouse lines were reported in previous studies. Ar*^f/Y^* strain with Exon 2 flanked by LoxP sites was obtained from and previously described by the laboratory of Dr. Guido Verhoeven^26^. To induce the Cre recombinase activity in nucleus, 2 doses of tamoxifen (3mg/dose in corn oil) were injected I.P. to 6-8 weeks old mice with interval of 48 hours.

All mouse genotypes were identified by PCR. The genotyping primers are listed in Supplementary Resource Table.

### Mouse prostate-tissue dissociation

The dissociation of mouse prostate tissues was previously described^24^. After the mice were sacrificed, prostate tissues were token off and sheared by ophthalmic tweezers and scissors. Prostate-tissue fragments were digested with 1mL 1X collagenase/hyaluronidase (STEMCELL, no. 07912) and 1 mL TrypLE (Gibco, no. 12605-028) separately for 30 minutes and 10 minutes at 37°C with gentle shaking, which contain 10 μM Y27632. TrypLE were neutralized in DMEM containing 10% FBS. The digested prostate cells were collected by centrifugation at 300 g for 3 minutes.

### Immunohistochemistry (IHC)

Mouse prostate tissues were collected, fixed in 4% PFA, dehydrated in ethanol, cleared with xylene, and embedded in paraffin. Paraffin embedding, sectioning, and H&E staining were performed by The Center of Comparative Medicine and Pathology (CCMP) Core Facility, the Molecular Cytology Core Facility at MSK (New York, NY) or Histeoserv Inc (Germantown, MD). Immunohistochemistry staining was performed on Ventana automatic stainer. Both HE and IHC sections were scanned using Mirax Digital Slide scanner. The catalog and dilution information of antibodies are in Supplementary Resource Table.

### Immunofluorescence (IF)

Indirect immunofluorescence was used on paraffin slides using signal amplification was conducted using Leica Bond BX staining system. Paraffin-embedded tissues were sectioned at 5 μm and baked at 58°C for 1 hr. Slides were loaded in Leica Bond and IF staining was performed as follows. Samples were pretreated with EDTA-based epitope retrieval ER2 solution (Leica, AR9640) for 20 mins. at 95°C. The 5-plex antibody staining and detection were conducted sequentially. For rabbit antibodies, Leica Bond Polymer anti-rabbit HRP was used, for the goat and chicken antibodies, a rabbit anti-goat, and a rabbit anti-chicken (Jackson ImmunoResearch303-006-003) secondary antibodies were used as linkers before the application of the Leica Bond Polymer anti-rabbit HRP. After that, Alexa Fluor tyramide signal amplification reagents (Life Technologies, B40953, B40958) or CF® dye tyramide conjugates (Biotium, 92172, 96053, 92174) were used for detection. After each round of IF staining, Epitope retrieval was performed for denaturation of primary and secondary antibodies before another primary antibody was applied. After the run was finished, slides were washed in PBS and incubated in 5 μg/ml 4’,6-diamidino-2-phenylindole (DAPI) (Sigma Aldrich) in PBS for 5 min, rinsed in PBS, and mounted in Mowiol 4–88 (Calbiochem). Slides were kept overnight at - 20°C before imaging. The catalog and dilution information of antibodies are in Supplementary Resource Table.

Direct immunofluorescence was used on frozen sections (**Figure 3**). The tissues were washed with precooled PBS three times and dehydrated by precooled 30% sucrose in PBS overnight at 4 °C. Dehydrated tissues were transferred into 30% optimal cutting temperature compound (OCT) in 30% sucrose for 1 h at 4 °C. Tissues were embedded in OCT and stored at−80 °C. Frozen samples were cut into 10 μm sections and stored at −20 °C. Sections were permeabilized in 1× PBS containing 0.1% Triton X-100 and blocked with 5% goat serum in 1× PBS containing 0.1% Triton X-100. Sections were then incubated with primary antibodies in 5% goat serum at room temperature for 2 h or overnight at 4 °C. The sections were washed with PBST (0.05% Tween 20 in 1× PBS) three times before incubation with secondary antibodies in 5% goat serum at room temperature in darkness for 1 h. The sections were washed with PBST three times, stained for nuclei with 4,6-diamidino-2-phenylindole (DAPI) (Sigma) and mounted with mounting medium.

For immunofluorescence of mouse organoids, organoids were resuspended in Cell Recovery Solution (Corning) to melt the Matrigel without disrupting the 3D cellular organization. Organoids were pelleted and embedded into fibrinogen and thrombin clots. These were transferred into embedding cassettes and fixed in 10% formalin. The paraffin sections were prepared at The Center for Translational Pathology of the Department of Pathology and Laboratory Medicine at Weill Cornell Medicine. Immunofluorescent staining was performed by using anti-Ck5 (BioLegend), anti-Ck8 (Abcam), and anti-FLAG primary antibodies (Novus Biologicals). Immunofluorescence images were collected as RGB composites, with DAPI in the blue channel and FLAG–Alexa 647 in the magenta channel. Images were analyzed in Python using scikit image. For each image, the DAPI channel was smoothed with a Gaussian filter (σ = 2 pixels), and nuclei were outlined using Otsu thresholding. Nuclear masks were cleaned by removing small spots (<100 pixels), filling holes, and eroding edges by 2 pixels to focus on the nuclear interior and limit signal from nearby cytoplasm. Individual nuclei were then labeled.

FLAG signal was measured from the red channel showing Alexa 647. For each nucleus, the average red intensity was recorded. To set a meaningful cutoff and avoid nonspecific staining, the magenta intensity measurements in negative-control nuclei (EV DMSO) were used to set the cutoff at the 99.5th percentile. This single cutoff was used for all images. Nuclei with average magenta intensity above this value were called FLAG-positive. Each sample’s total nuclei, FLAG-positive nuclei, and percent FLAG-positive cells were recorded. All analysis settings were kept constant across conditions.

### Transmission electron Microscopy

Prostate tissue from 10-week-old, 2-week tamoxifen-treated TY, TYA, and TY-castrate mice were dissected. The proximal and distal portions of the anterior lobes were further dissected and fixed in 2% glutaraldehyde, 4% PFA, 2 mM CaCl2 in 0.1 M sodium cacodylate buffer, pH 7.2, for > 1 hour, postfixed in 1% osmium tetroxide, dehydrated in an acetone series and processed for Eponate 12 (Ted Pella, Inc) embedding. Ultrathin sections (65 nm) were cut and post-stained with uranyl acetate/lead citrate and imaged in a Tecnai 12 electron microscope (FEI, Hillsboro, OR) operating at 120kV equipped with an AMT BioSprint29 digital camera (Woburn, MA).

### Mouse organoid culture

To generate organoids with inducible dnAP-1 expression, prostate from B6 mice were harvested at 1-2 months of age and processed and grown as 3D Matrigel culture as previously described with the exception that the organoid media contained 5 ng/mL EGF^48^. Doxycycline (Dox)-inducible Acidic-FOS(A-FOS) system, referred to as dnAP-1, was engineered via pENTR/directional TOPO cloning of the FLAG epitope-tagged A-FOS construct2 from a CMV500 backbone (Addgene #33353) into the doxycycline-inducible pCW57.1 vector (Addgene #41393). Plasmid sequences were verified by whole-plasmid sequencing (Plasmidsaurus). Polyclonal populations were derived via lentivirus-mediated transduction, followed by puromycin selection and immunoblotting using a FLAG antibody to verify A-FOS induction with 1 μg/mL Dox treatment for 24 hours.

To characterize the role of dnAP-1 on organoid histology and gene expression, the engineered organoids were seeded into Matrigel without EGF. Organoids were seeded at a starting density of 500 cells per dome and cultured in an Incucyte S3 for 6 days in media containing 1 µg/mL DOX (for doxycycline conditions), 10 nM DHT (for DHT conditions). Live-cell images were captured every 6 hours at 4x magnification. Organoid size and area were measured and quantified using OrganoID. Statistical analyses was performed using two-tailed non-parametric Welch’s t-test for a total of four biological replicates.

To assay organoid formation of the wild type and Ar knockout luminal cells, we FACS sorted Epcam+,CD49f^low^ luminal cells from TY and TYA mouse prostates 2 weeks after tamoxifen injection, and Trop2^+^ and Trop2^-^ luminal cells were separated and seeded 500 cells per Matrigel dome and cultured with organoid medium containing 1 nM DHT for 8 days, then counted number of organoids under microscope.

### Immunoblot

Whole-cell protein lysates were prepared after digestion of Matrigel with TrypLE Express Enzyme (1X), no phenol red (GIBCO). Pelleted cells were washed in PBS and lysed in RIPA buffer supplemented with protease and phosphatase inhibitors. Proteins were quantified by BCA assay and separated on TGX 4%–15% Protein Gels. The catalog and dilution information of antibodies are in Supplementary Resource Table.

### Flow cytometry assay

Freshly dissociated prostate cells were resuspended by 200 μL FACS buffer (1x PBS containing primocin, 10 mM HEPES, and 1% BSA, supplemented with 1 nM Y27632) and stained with antibodies for 30 minutes on ice (Supplementary table 1). And then the stained prostate cells were washed with 1 mL FACS buffer and passed through a 40 μm filter. Prostate samples were analyzed by LSRFortessa (BD) and sorted using FACS Aria III (BD).

### Single cell RNA sequencing (scRNAseq)

*Tmprss2-CreER^T2^-IRES-nlsEGFP*; *EYFP* and *Tmprss2-CreER^T2^-IRES-nlsEGFP*; *EYFP*; *Ar*^f/Y^ mice were euthanized after tamoxifen treatment. After euthanasia, the prostates were dissected out and minced with scalpel and then processed for 1h digestion with collagenase/hyaluronidase (#07912, STEMCELL Technologies) and 15min digestion with TrypLE (#12605010, Gibco). Dissociated prostate cells were sorted out by flow cytometry as DAPI negative. The 3 replicates of each condition were labeled with TotalSeq™ anti-mouse Hashtag antibodies (B0301/B0302/B0303, #155831, BioLegend) separately and mixed as one sample. For each mixture, 10,000 cells were directly processed with 10X genomics Chromium Single Cell 3’ GEM, Library & Gel Bead Kit v3 according to manufacturer’s specifications. For each sample, 200 million reads were acquired on NovaSeq platform S4 flow cell.

Reads obtained from the 10x Genomics scRNAseq platform were mapped to mouse genome (mm10) including the transgenes (*CreER^T2^* and EYFP) using the Cell Ranger multi software (10X Genomics). True cells were distinguished from empty droplets using scCB2 package^49^. Down-stream analysis and figure plotting are done by Scanpy (version 1.9.8)^50^. The levels of mitochondrial reads and numbers of unique molecular identifiers (UMIs) were similar among the samples, which indicates that there were no systematic biases in the libraries from mice with different genotypes. Cells were removed if they expressed fewer than 200 unique genes, less than 2,000 total counts, more than 50,000 total counts or greater than 20% mitochondrial reads. Genes detected in less than 10 cells and all mitochondrial genes were removed for subsequent analyses. Putative doublets were detected and filtered out using the doublet detection package (http://doi.org/10.5281/zenodo.2678041). The average gene detection in each cell type was similar among the samples. Combining samples in the entire cohort of Control and ARKO groups yielded a filtered count matrix of 92,938 cells by 19,448 genes, with a median of 12984 counts and a median of 3135 genes per cell. The count matrix was normalized by log2(10K+1) for calculating the top 2000 highly variable genes using Scanpy. The count matrix was further scaled to mean 0 and standard deviation 1 for PCA analysis and UMAP dimension reduction (https://arxiv.org/abs/1802.03426), and Leiden clustering^51^. Principal Component Analysis (PCA) was performed on the 1,000 most variable genes and the top 50 principal components (PCs) retained 44% variance explained. Marker genes for each cluster were found with Scanpy. Cell types were determined using a combination of marker genes identified from the literature and gene ontology for cell types using the web-based tool Panglao DB (https://panglaodb.se/).

The mean expression of each cluster and each sample were generated with Scanpy. GSEA analyses were performed using JAVA GSEA 4.1.0 program, using curated gene sets (C2) and Hallmark gene sets (H) from Molecular Signatures Database. Imputation was performed using MAGIC (Markov affinity-based graph imputation of cells) package^52^.

### Bulk RNA sequencing

For bulk RNA sequencing, total RNA was extracted from FACS sorted EYFP^+^,Epcam^+^,Cd24a^hi^ prostate cells with TRIzol reagent following the manufacture’s instruction (#15596026, Invitrogen). RNAseq libraries were prepared using standard Illumina PolyA library preparation protocol. Next generation sequencing was performed by MSKCC genomics core facility on Illunima HiSeq2500 platform with paired end of 50bp for 30-40 million reads. The sequence data were mapped to the mouse genome (mm9) using STAR^53^. Gene expressions were normalized with total mapped reads among different groups. Differentially expressed genes were determined as absolute fold change >2, p<0.05 in R with DESeq2 (1.46.0).

For bulk RNA sequencing of prostate organoids, prior to seeding, 1ug/mL doxycycline was premixed into the Matrigel-cell suspension for all DOX conditions. Murine Organoids were grown for 4 days in media containing 1ug/mL DOX (for dox conditions), 1 nM DHT, and then for 24 hr without DHT and EGF. Next, cells were stimulated with 1ug/mL DOX (for dox conditions), 10nM DHT without EGF for 24 hr and harvested for RNA-seq in biological triplicates. Total RNA was extracted using RNeasy Plus Mini Kit. DNA was removed via column-based extraction per manufacturer’s protocol, followed with Nanodrop quantification, and quality check using Agilent Bioanalyzer RNA 6000 Pico. Samples with RNA integrity number > 9.5 were used for library preparation (NEB Ultra II Directional RNA Library Prep (plus Poly A isolation module) at the Weill Cornell Medicine Genomics Core. Libraries were pair-end sequenced on a Novaseq X Plus (Illumina) in 1.5B Flow Cell mode to generate 100 bp reads. Data was processed using nf-core collection of workflows (v3.10). The pipeline was executed with Nextflow (v23.04.3) with the following command: nextflow run nf-core/rnaseq-r 3.12.0 --input samplesheet.csv --genome GRCm38-profile singularity. Briefly, raw FASTQ files were aligned to the GRCm38 (mm10) reference genome and quantified using salmon (v 1.10.0). Differential gene expressions were performed in R with DESeq2 (1.46.0).

GSEA analyses were performed using JAVA GSEA 4.1.0 program, using curated gene sets (C2) and Hallmark gene sets (H) from Molecular Signatures Database. Genes sets of luminal distal and proximal mouse prostate cells were derived from prior work^10^. Plots were generated using ggplot2 or GraphPad Prism.

### Bulk ATAC sequencing

Transposase-accessible chromatin sequencing (ATAC-seq) of FACS sorted EYFP^+^,Epcam^+^,Cd24a^hi^ cells was performed as previously described^24^. For each sample, 5,000 viable cells were processed for nuclei isolation and transposase treatment. The digestions were carried out for 30 min at 37. The library preparation and next generation sequencing were performed by MSKCC Genomics Core Facility. For each sample, 40-50 million paired-end reads were sequenced on Illunima HiSeq2500 platform. The sequence data were processed for adapter trimming and aligned to mouse genome (mm10) with bowtie 2^54^. Peaks were called using MACS2^55^, the command is “macs2 callpeak-t bam-f BAMPE-g mm --call-summits”. Only the overlapped peaks from 2 replicates were kept for downstream analysis. All of the reproducible peaks (overlapped peaks in 2 biological replicates) form all samples TY, TYA, and TY-cast were merged and annotated by Homer (annotatePeaks.pl)^56^, then the enhancer and promoter peaks were separated based on Homer annotation. The reads in the peaks were counted by featureCounts^57^. The log fold changes (LFC) and adjusted p values (padj) were calculated by DESeq2 package^58^.

### Quantification and statistics of mouse prostate cell numbers

The Trop2, EYFP, Ar, Ki67, EdU, Ck8, Ck5, Trp63, tdTomato, ZsGreen positive cell numbers were quantified and averaged on 3 representative areas on IF or IHC images from distal or proximal anterior prostate of each mouse. The tissue area was quantified on H&E-stained slides using Fiji ImageJ software. At least three mice were used for each condition. Significance was evaluated by two tails unpaired Student’s t-test. For organoid formation assay, the data were obtained from at least three independent experiments.

## Data and materials availability

The accession numbers for bulk RNA-seq, single cell RNA-seq and ATAC-seq are as follows: RNA-seq data (primary luminal cells: GSE306685, organoid cells: GSE314378), scRNA-seq data (GSE306891), ATAC-seq data (GSE306686).

## Supporting information

Supplementary Resource Tables

## Acknowledgements

We would like to thank the MSK Molecular Cytology Core for immunohistochemistry and immunofluorescence staining, the MSK Single-Cell Analytics Innovation Lab (SAIL) and the MSK Genomics Core Facility for advice, the Rockefeller University Electron Microscopy Resource Center.

## Author contributions

Y.C., P.C., D.G. and D.L. conceived and designed the experimental approach. D.L., N.W. and W.G. performed most experiments. J.O. performed dnAP-1 experiments. D.L. and Y.C. contributed to the computational and statistical analysis. W.H.C., D.S., S.C., H.Z., U.I.C., C.K.W., V.R.C., H.W., W.K., N.F. and C.E.B. helped with the experiments and provided technical support. H.A.P. and A.S. performed EM imaging. A.G. performed pathology on mouse samples. Y.C., P.C., D.G. and D.L. prepared the manuscript as the senior authors.

## Supplementary Figure Legends

**Figure S1.**
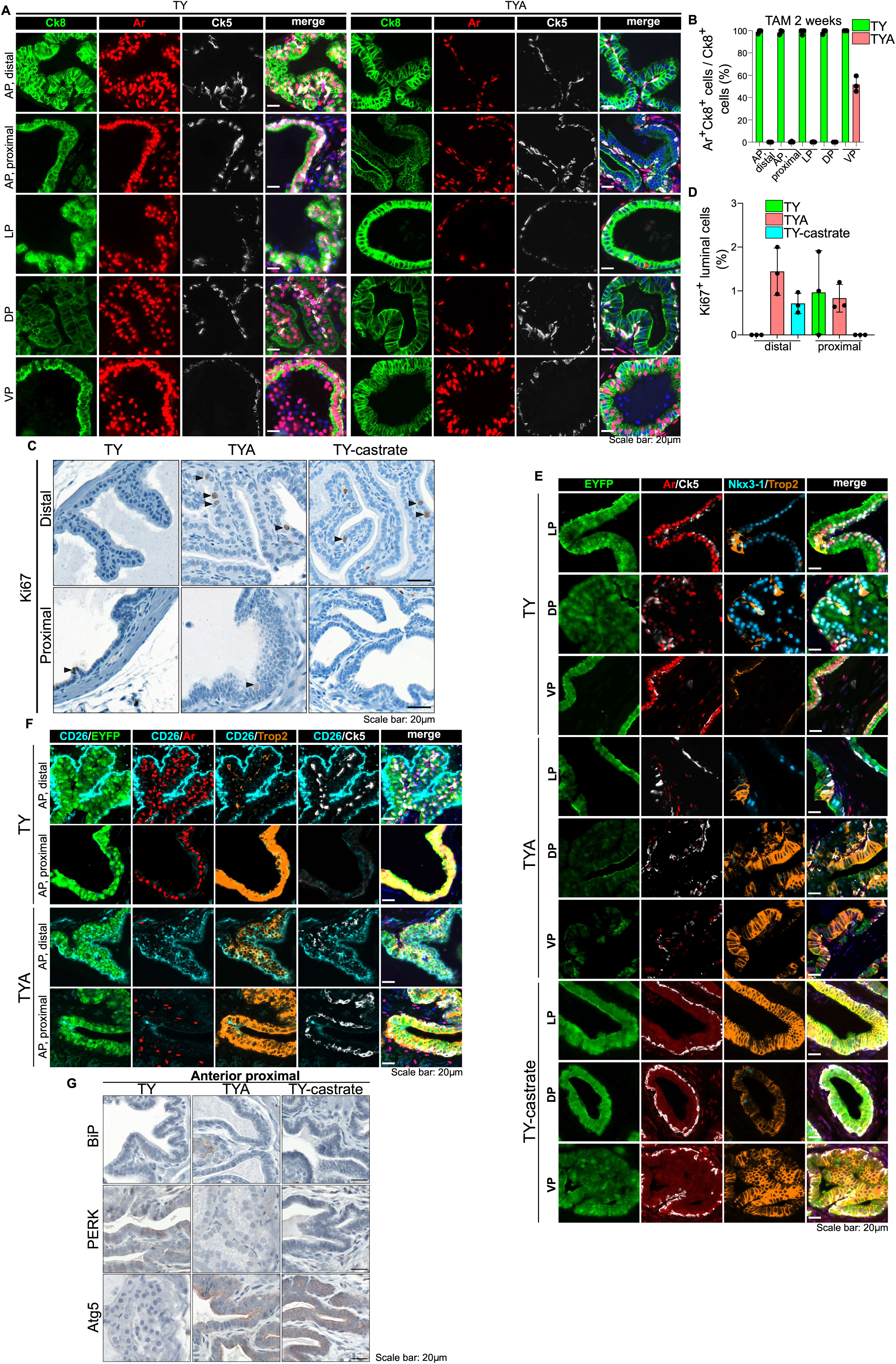
Immunohistochemistry characterization of luminal cell specific Ar deletion or castration. (**A**) Representative multiplex IF of indicated proteins in distal and proximal regions of anterior prostates (AP), distal lateral (LP), distal dorsal (DP), and distal ventral prostates (VP) in TY and TYA mice 2 weeks after tamoxifen treatment. (**B**) Quantification (percentage of Ar^+^Ck8^+^/Ck8^+^ cells) of different prostate lobes from IF staining in panel A (n=3 mice). (**C**) Representative Ki67 IHC in distal and proximal regions of anterior prostates in TY, TYA and TY-castrate mice. (**D**) Quantification (percentage of Ki67^+^ luminal cells) from IHC staining in panel C. (**E**) Representative multiplex IF of indicated proteins are shown in TY, TYA and TY-castrate mice. (**F**) Representative multiplex IF of noted proteins are shown in TY and TYA mice. (**G**) Representative IHC staining of ER-associated proteins (BiP, PERK) and autophagy-related protein Atg5 are shown in proximal anterior prostates in TY, TYA and TY-castrate mice. Scale bar: 20mm

**Figure S2.**
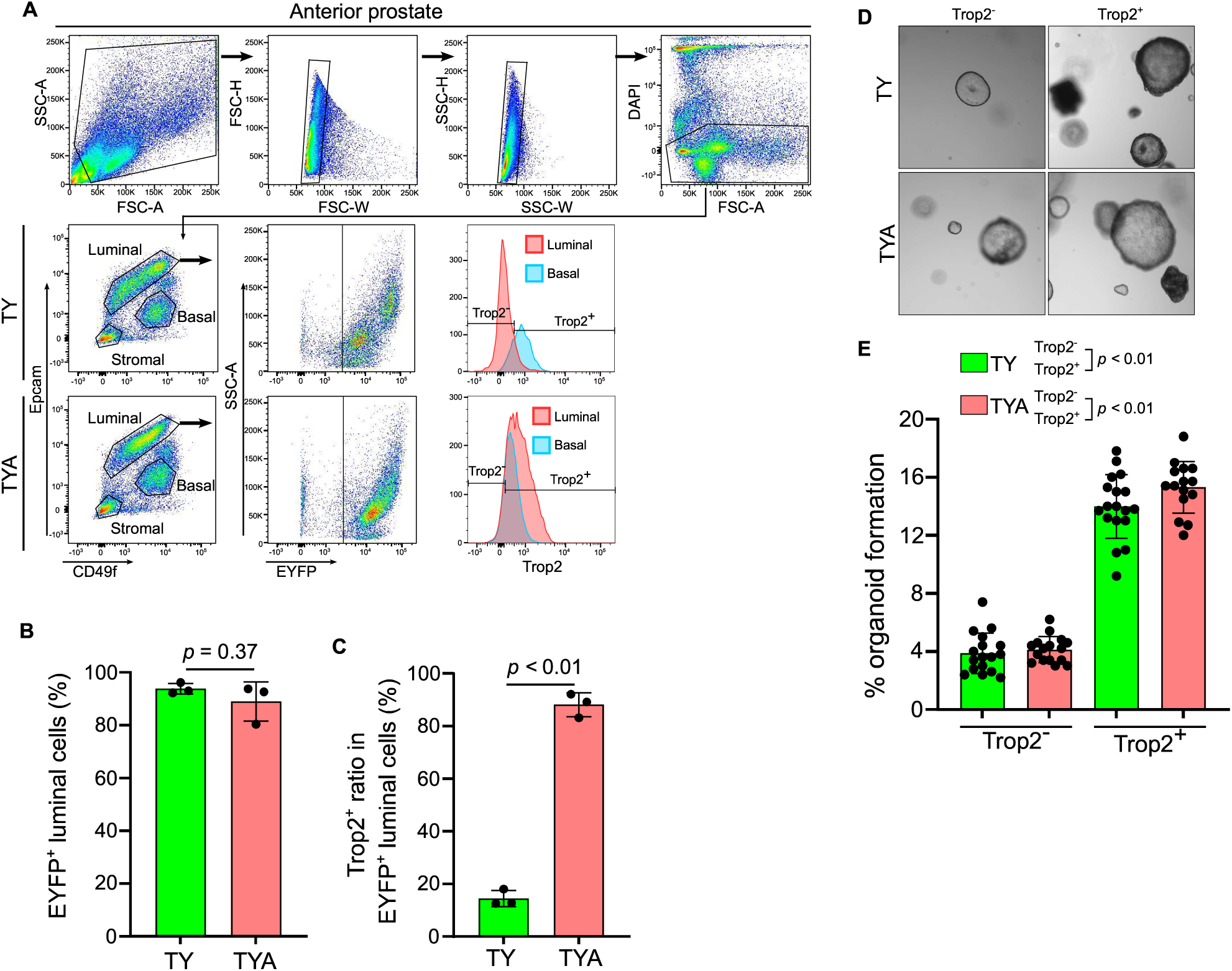
Ar loss-induced Trop2^+^ luminal cells have higher organoid formation capacity. (**A**) FACS gating strategy to identify Epcam^Lo^, CD49f^Hi^ basal cells and Epcam^Hi^, CD49f^Lo^ luminal cells. The proportion of Trop2^+^ luminal cells threshold is based on basal cells. (**B-C**) FACS-based quantification of EYFP and Trop2 positivity in TY and TYA luminal cells. (**D**) Representative images of organoids formed from Trop2^-^ and Trop2^+^ luminal cells. (**E**) Quantification (number of organoids per 100 luminal cells) from panel D. Data are presented as mean ± s.d. and analyzed with two-tailed *t*-test. Scale bar, 20 mm.

**Figure S3.**
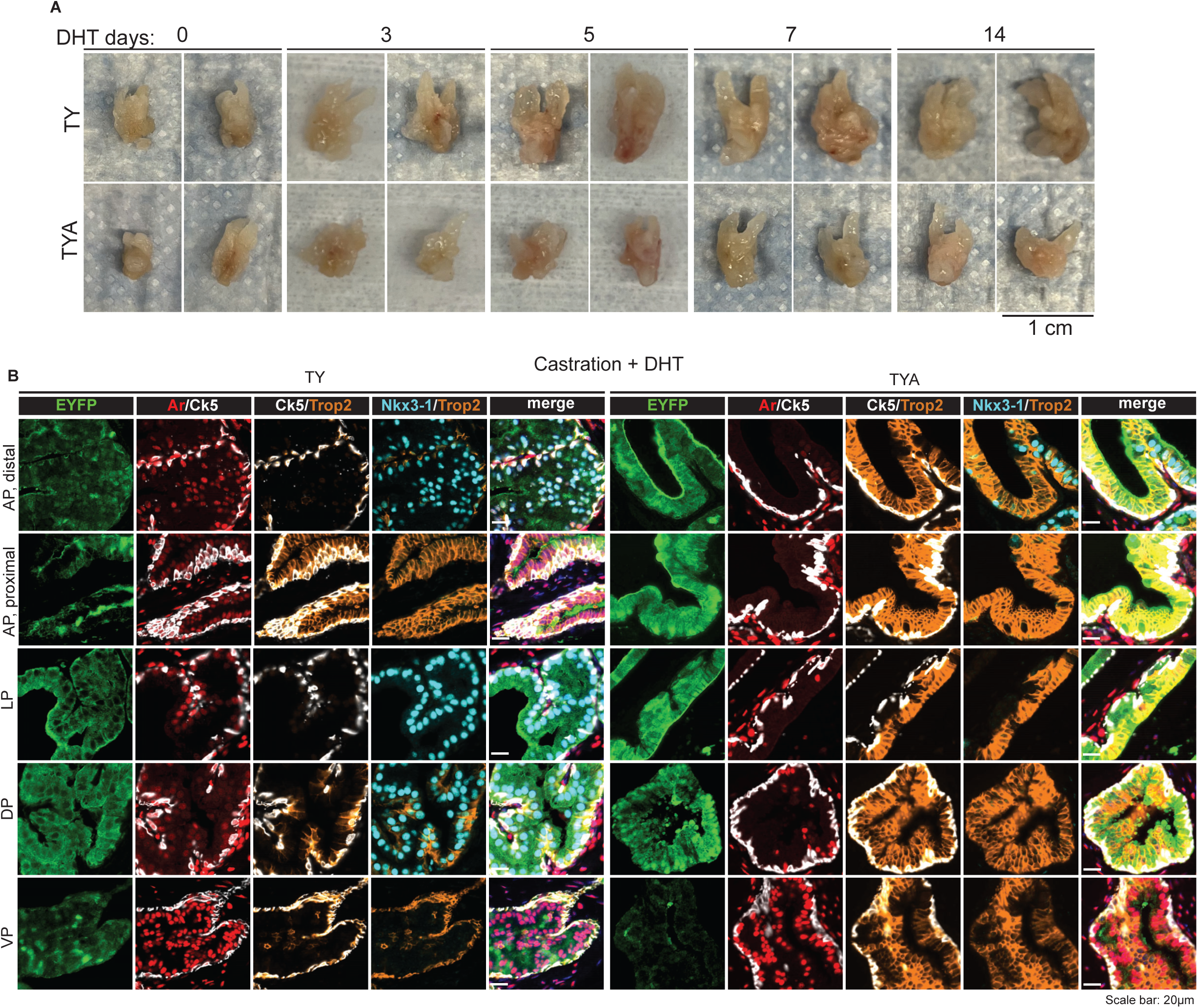
Luminal Ar leads to defective prostate regeneration (**A**) Images of TY and TYA prostates at indicated time points after DHT treatment 2 weeks after castration. (**B**) Representative multiplex immunofluorescence (IF) of indicated proteins in distal and proximal regions of anterior prostates (AP), distal lateral (LP), distal dorsal (DP), and distal ventral prostates (VP) in TY and TYA mice 2 weeks after castration followed by 2 weeks DHT treatment.

**Figure S4.**
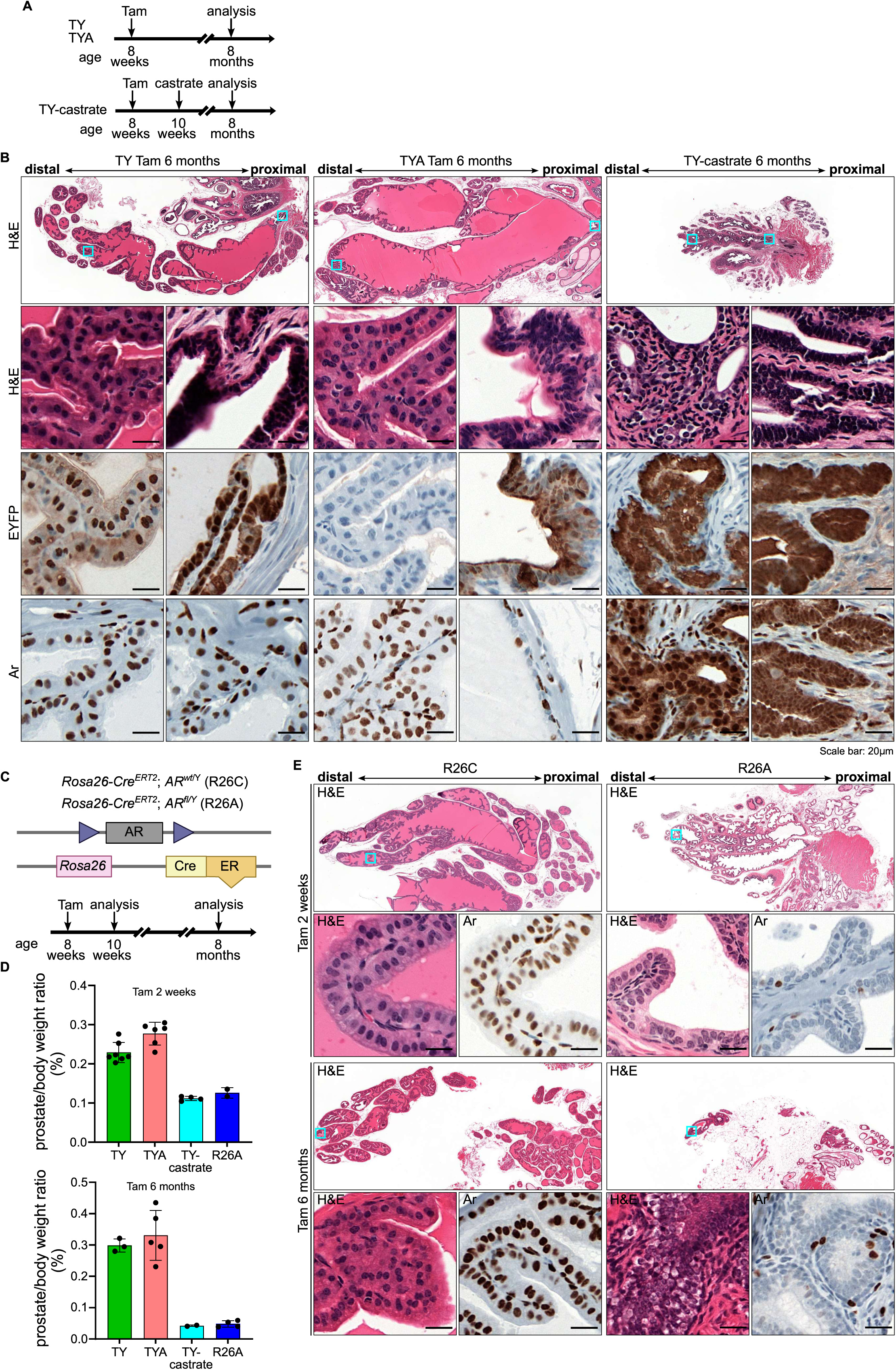
Ar-deleted luminal cells are eliminated over time. (**A**) Schematic of the experimental strategy. (**B**) Representative histological changes are shown in distal and proximal anterior prostates of TY, TYA and TY-castrate mice 6 months after tamoxifen injection (TY and TYA) or 6 months after castration (TY-castrate). (**C**) Schematic of the targeting strategy to generate *Rosa26-CreER^T2^;Ar^wt/Y^* (R26C) and *Rosa26-CreER^T2^;Ar^f/Y^* (R26A) mice. (**D**) The mouse body and prostate weights are measured 2 weeks and 6 months after tamoxifen injection (TY, TYA and R26A), or 2 weeks and 6 months after castration (TY-castrate). Data is presented as mean ± s.d.. (**E**) Representative histological changes are shown in distal anterior prostates of R26C and R26A mice 2 weeks and 6 months after tamoxifen injection. Ar expression is shown with IHC staining.

**Figure S5.**
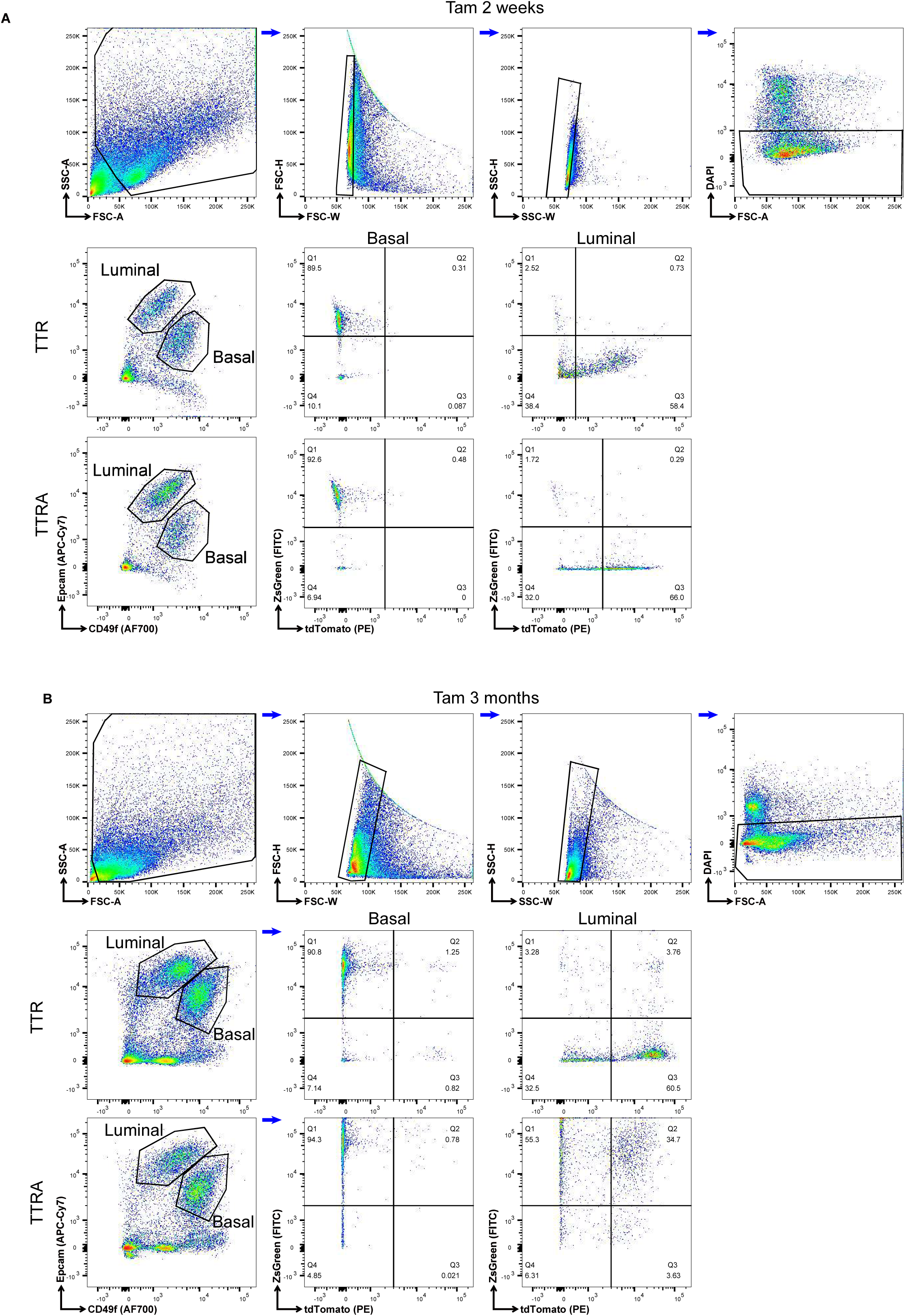
FACS quantification of cell populations of TTR and TTRA mice (**A**) Quantification of ZsGreen and tdTomato positivity in luminal and basal cells from TTR and TTRA mice 2 weeks after tamoxifen injection using FACS. (**B**) Quantification of ZsGreen and tdTomato positivity in luminal and basal cells from TTR and TTRA mice 3 months after tamoxifen injection.

**Figure S6.**
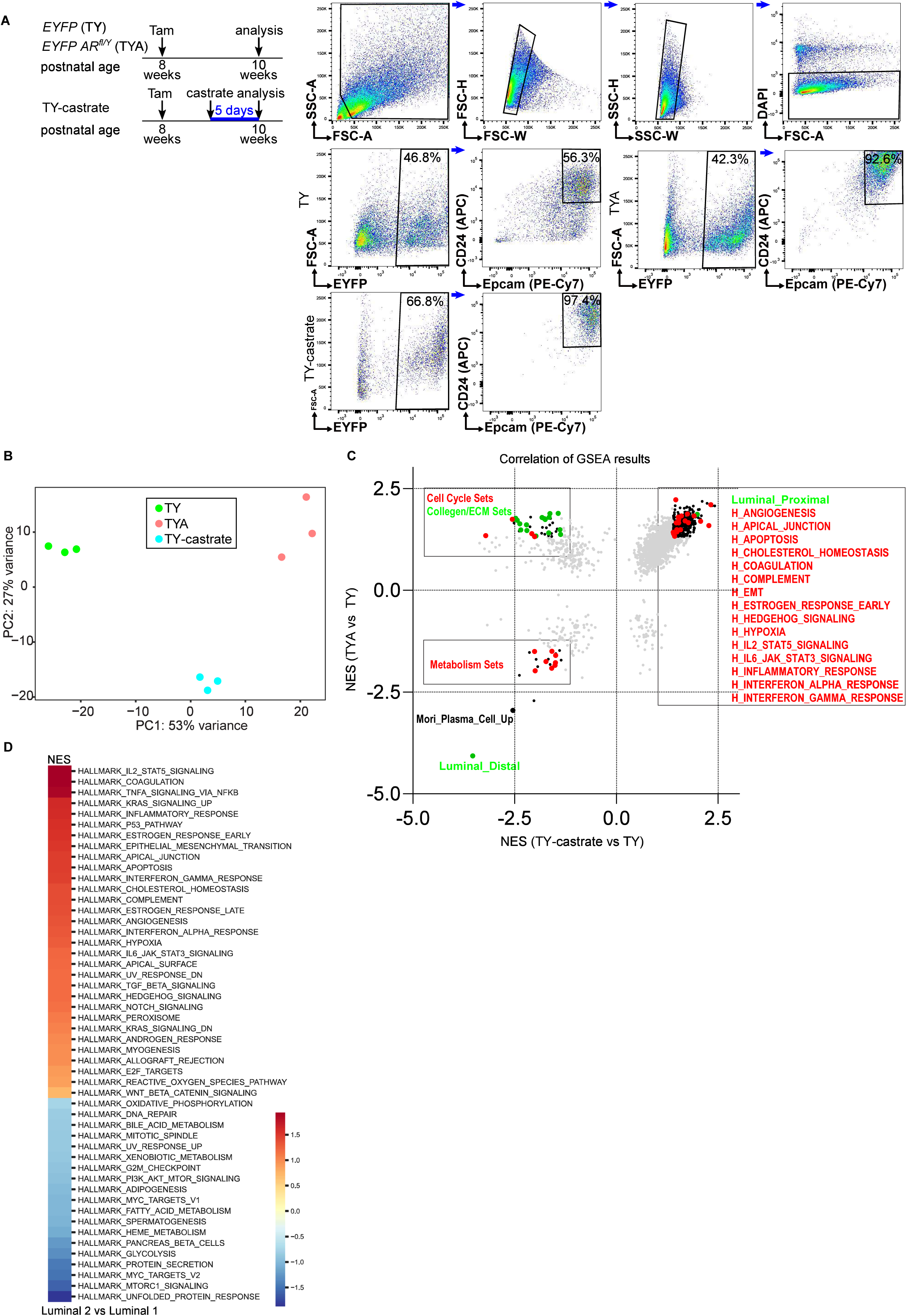
Transcriptomic survey shows Ar knockout luminal cells specifically increase cell cycle and extracellular matrix gene sets compared with castrated luminal cells (**A**) EYFP^+^/Epcam^+^/CD24^high^ luminal cells from TY, TYA and TY-castrate mice are sorted for bulk RNA sequencing. (**B**) Principal components analysis (PCA) of bulk RNA sequencing data of sorted TY, TYA and TY-castrate luminal cells. (**C**) GSEA normalized enrichment scores of enriched gene sets in TYA compared with TY, TY-castrate compared with TY. (**D**) GSEA normalized enrichment scores of Hallmark genes sets in Luminal 2 compared with Luminal 1 in intact TY mice.

**Figure S7.**
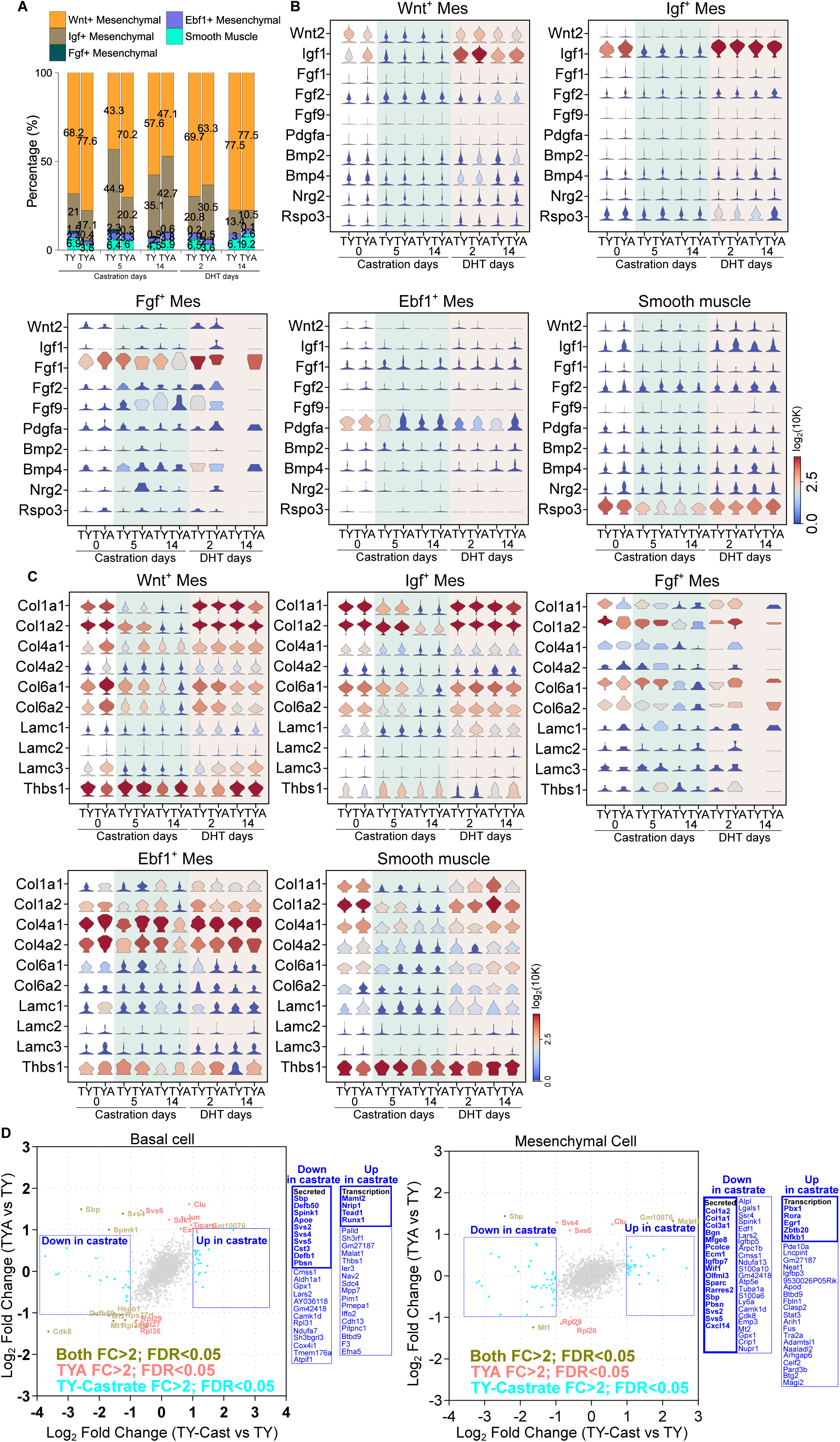
Changes in markers of mesenchymal cells after castration and DHT treatment in TY and TYA mice. (**A**) The percentage of different mesenchymal cell types are quantified in single-cell RNA-seq data. (**B**) Violin plots of mesenchymal cell expressed growth factors. The color in the violin plots indicates the median expression level of genes. (**C**) Violin plots of mesenchymal cell expressed extracellular matrix genes. The color in the violin plots indicates the median expression level of genes. (**D**) Scatter plot of differentially expressed genes of TYA compared with TY, TY-castrate compared with TY by single cell RNA-seq of basal and mesenchymal cells.

**Figure S8.**
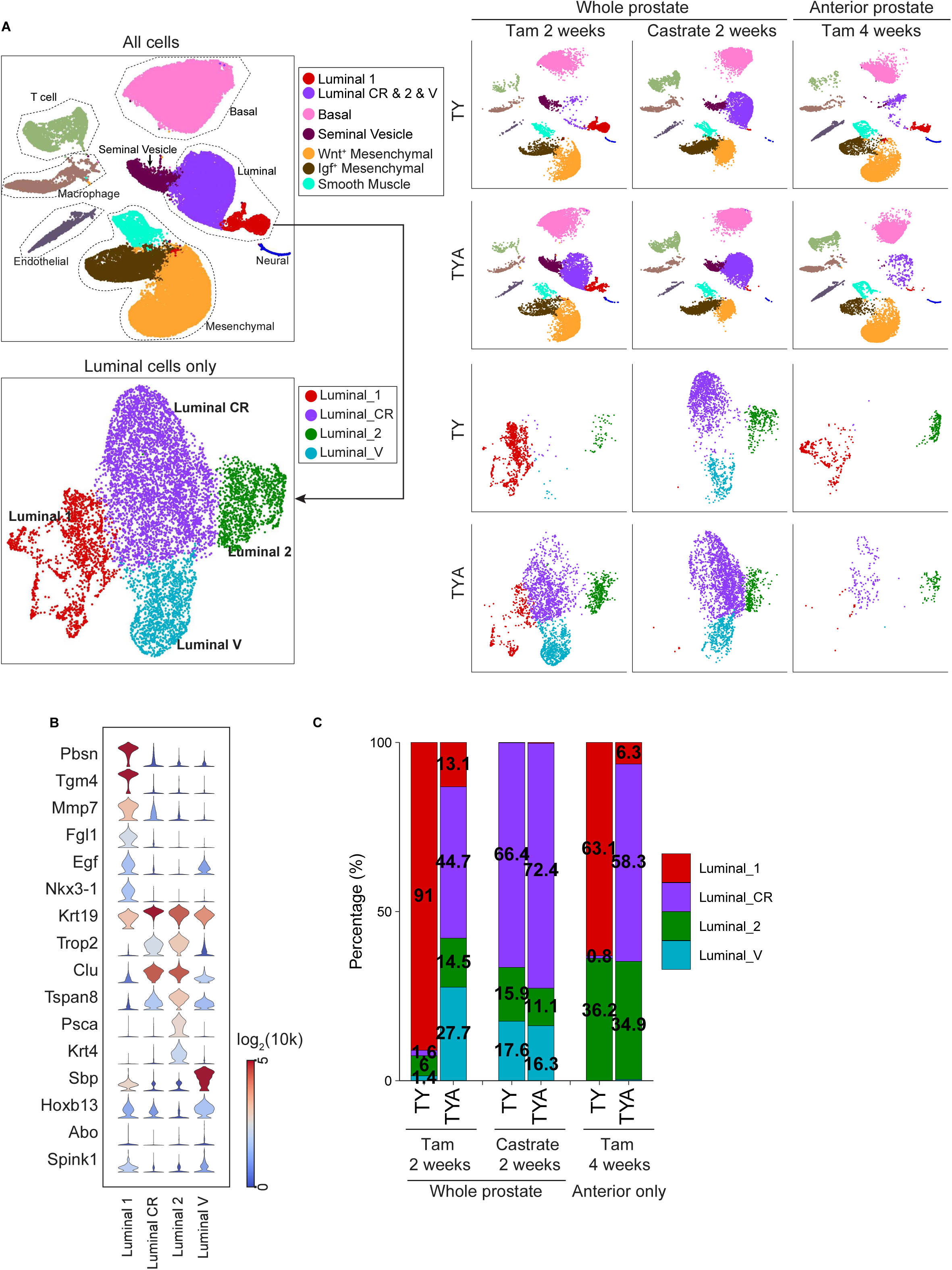
Independent validation dataset of single-cell RNA-seq of anterior prostate (**A**) Single-cell census of the anterior prostates, which are integrated with whole prostate data. Uniform manifold approximation and projection (UMAP) of single-cell RNA-seq profiles colored by Leiden clustering of subsets are shown. (**B**) Violin plots of Luminal 1, Luminal CR, Luminal 2 and Luminal V markers. The color in the violin plots indicates the median normalized expression level of genes. (**C**) The percentage of different Luminal cells are quantified in single-cell RNA-seq data.

**Figure S9.**
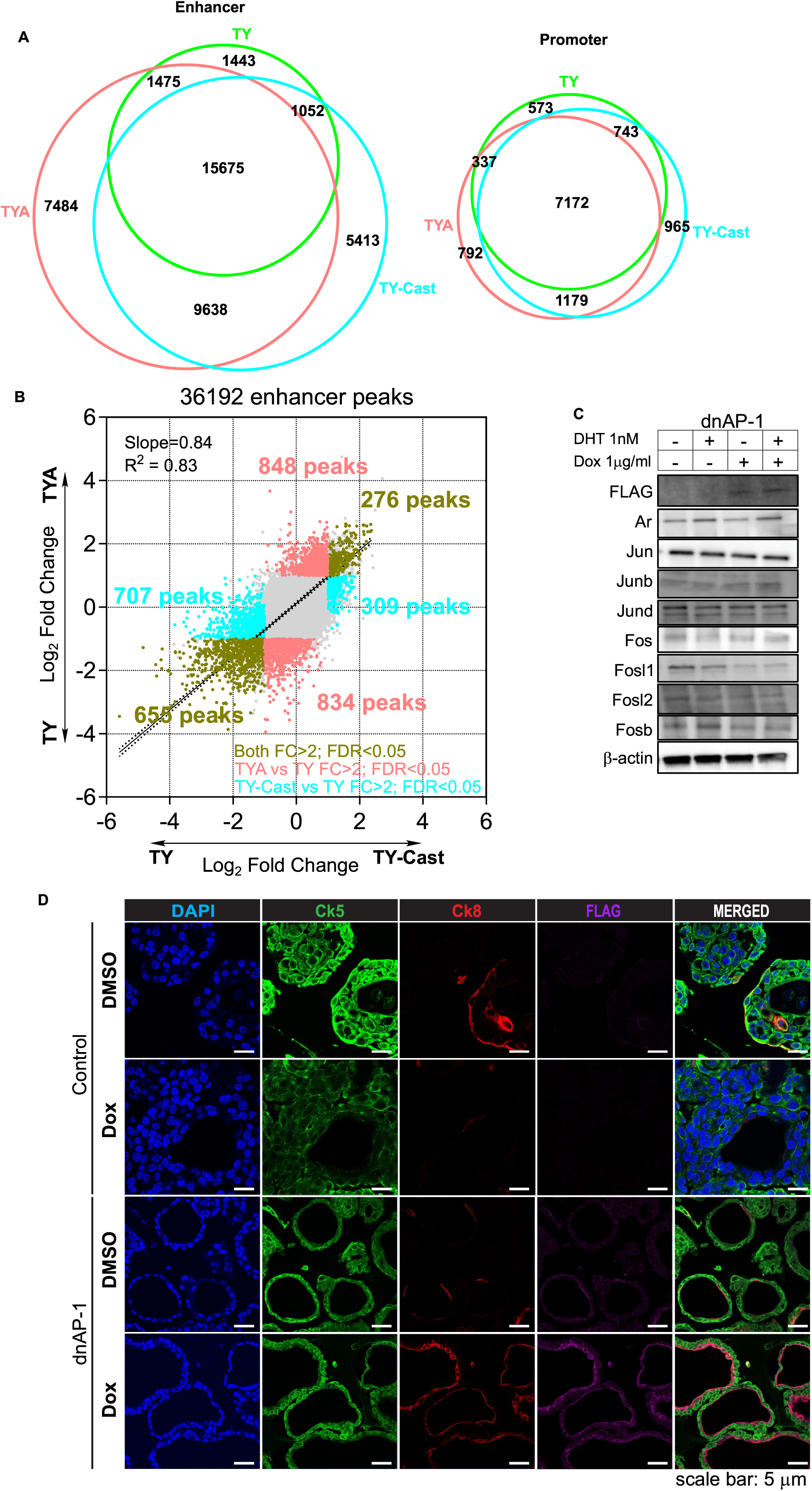
AP-1 inhibition promotes luminal lineage specification *in vitro* (**A**) Venn diagram of the overlap of enhancer and promoter ATAC peaks of sorted luminal cells from TY, TYA, and TY-castrate mice. (**B**) Scatter plot and correlation analysis of differential ATAC peaks of TYA compared with TY, TY-castrate compared with TY using sorted luminal cells. (**C**) Immunoblot of FLAG-tagged A-Fos and indicated proteins after 24 hours treatment with and without 1 mg/ml Dox and 1 nM DHT. (**D**) Immunofluorescence of prostate epithelial lineage markers in control and doxycycline-inducible dnAP-1 organoids 24 hours after Dox and or DMSO treatment.

**Figure S10.**
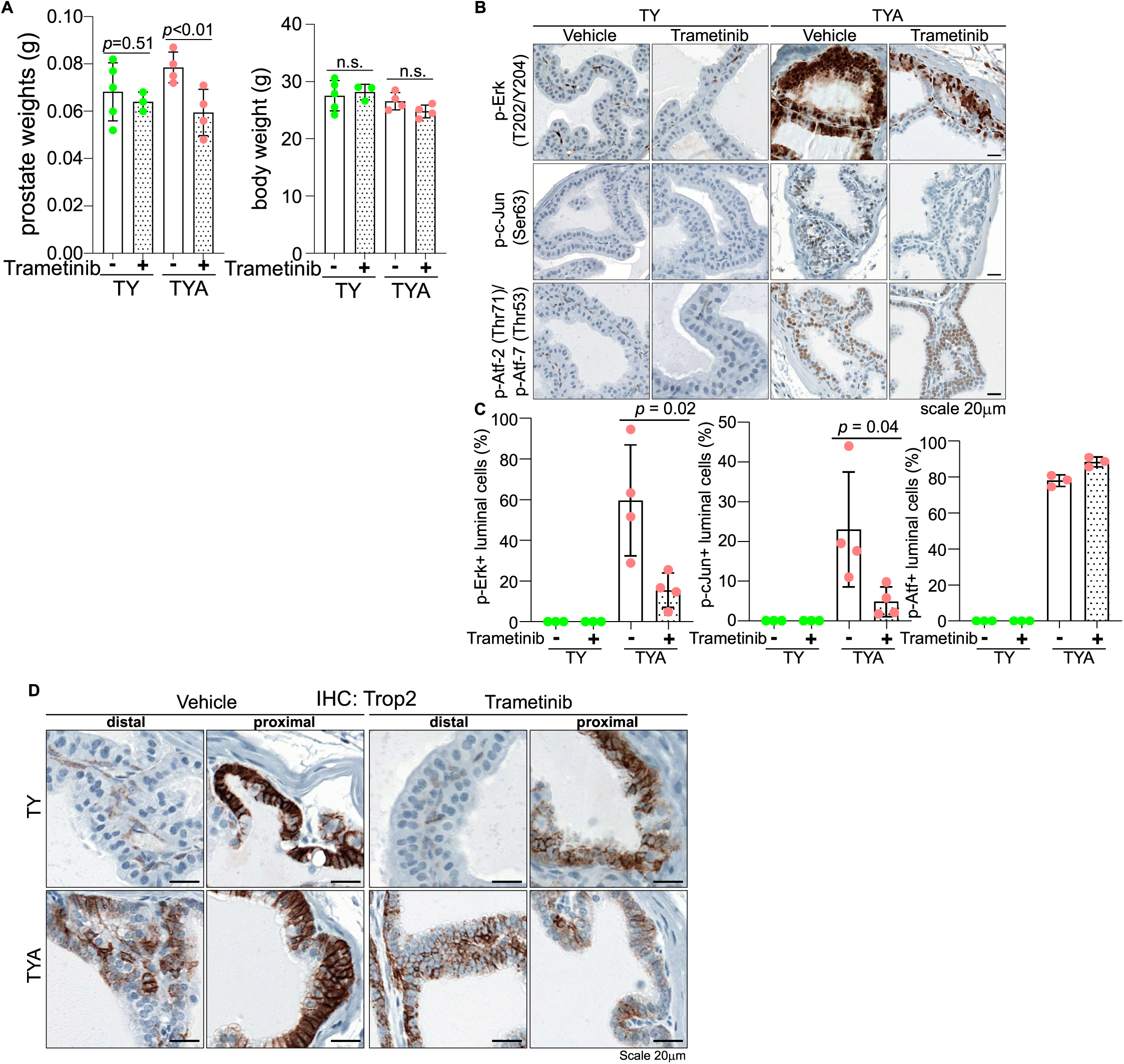
Effect of MEK inhibition in wild-type and Ar-knockout luminal cells (**A**) The mouse prostate and body weights are measured 1 week after trametinib treatment (1mg/kg, daily oral gavage). Data is presented as mean ± s.d. (**B**) Representative IHC staining of indicated proteins in distal anterior prostates in TY and TYA mice with or without trametinib treatment are shown. (**C**) Quantification of IHC positive cells ratio in panel B. (**D**) Representative Trop2 IHC in distal and proximal regions of anterior prostates in TY and TYA mice after trametinib treatment.

## References

1. Cunha, G.R. Mesenchymal-epithelial interactions during androgen-induced development of the prostate. Progress in clinical and biological research 171, 15–24 (1985).

2. Wu, C.T. et al. Increased prostate cell proliferation and loss of cell differentiation in mice lacking prostate epithelial androgen receptor. Proc Natl Acad Sci U S A 104, 12679–12684 (2007).

3. Simanainen, U. et al. Disruption of prostate epithelial androgen receptor impedes prostate lobe-specific growth and function. Endocrinology 148, 2264–2272 (2007).

4. Yu, S. et al. Altered prostate epithelial development in mice lacking the androgen receptor in stromal fibroblasts. Prostate 72, 437–449 (2012).

5. Zhang, B. et al. Non-Cell-Autonomous Regulation of Prostate Epithelial Homeostasis by Androgen Receptor. Mol Cell 63, 976–989 (2016).

6. Xie, Q. et al. Dissecting cell-type-specific roles of androgen receptor in prostate homeostasis and regeneration through lineage tracing. Nat Commun 8, 14284 (2017).

7. Chua, C.W. et al. Differential requirements of androgen receptor in luminal progenitors during prostate regeneration and tumor initiation. Elife 7 (2018).

8. Lee, D.H. et al. Androgen action in cell fate and communication during prostate development at single-cell resolution. Development 148 (2021).

9. Henry, G.H. et al. A Cellular Anatomy of the Normal Adult Human Prostate and Prostatic Urethra. Cell Rep 25, 3530–3542 e3535 (2018).

10. Aparicio, L. et al. Meta-analyses of mouse and human prostate single-cell transcriptomes reveal widespread epithelial plasticity in tissue regression, regeneration, and cancer. Genome Med 17, 5 (2025).

11. Karthaus, W.R. et al. Regenerative potential of prostate luminal cells revealed by single-cell analysis. Science 368, 497–505 (2020).

12. Guo, W. et al. Single-cell transcriptomics identifies a distinct luminal progenitor cell type in distal prostate invagination tips. Nat Genet 52, 908–918 (2020).

13. Crowley, L. et al. A single-cell atlas of the mouse and human prostate reveals heterogeneity and conservation of epithelial progenitors. Elife 9 (2020).

14. Mevel, R. et al. RUNX1 marks a luminal castration-resistant lineage established at the onset of prostate development. Elife 9 (2020).

15. Kirk, J.S. et al. Integrated single-cell analysis defines the epigenetic basis of castration-resistant prostate luminal cells. Cell Stem Cell 31, 1203–1221 e1207 (2024).

16. Song, H. et al. Single-cell analysis of human primary prostate cancer reveals the heterogeneity of tumor-associated epithelial cell states. Nat Commun 13, 141 (2022).

17. Cancer Genome Atlas Research, N. The Molecular Taxonomy of Primary Prostate Cancer. Cell 163, 1011–1025 (2015).

18. Huggins, C. & Hodges, C.V. Studies on Prostatic Cancer. I. The Effect of Castration, of Estrogen and of Androgen Injection on Serum Phosphatases in Metastatic Carcinoma of the Prostate. Cancer Research 1, 293–297 (1941).

19. Watson, P.A., Arora, V.K. & Sawyers, C.L. Emerging mechanisms of resistance to androgen receptor inhibitors in prostate cancer. Nat Rev Cancer 15, 701–711 (2015).

20. Pomerantz, M.M. et al. The androgen receptor cistrome is extensively reprogrammed in human prostate tumorigenesis. Nat Genet 47, 1346–1351 (2015).

21. Grbesa, I. et al. Reshaping of the androgen-driven chromatin landscape in normal prostate cells by early cancer drivers and effect on therapeutic sensitivity. Cell Rep 36, 109625 (2021).

22. Chen, X. et al. Canonical androgen response element motifs are tumor suppressive regulatory elements in the prostate. Nat Commun 15, 10675 (2024).

23. Gao, D. et al. A Tmprss2-CreERT2 Knock-In Mouse Model for Cancer Genetic Studies on Prostate and Colon. PLoS One 11, e0161084 (2016).

24. Li, D. et al. ETV4 mediates dosage-dependent prostate tumor initiation and cooperates with p53 loss to generate prostate cancer. Sci Adv 9, eadc9446 (2023).

25. Madisen, L. et al. A robust and high-throughput Cre reporting and characterization system for the whole mouse brain. Nat Neurosci 13, 133–140 (2010).

26. De Gendt, K. et al. A Sertoli cell-selective knockout of the androgen receptor causes spermatogenic arrest in meiosis. Proc Natl Acad Sci U S A 101, 1327–1332 (2004).

27. Munro, S. & Pelham, H.R. An Hsp70-like protein in the ER: identity with the 78 kd glucose-regulated protein and immunoglobulin heavy chain binding protein. Cell 46, 291–300 (1986).

28. Yousefi, S. et al. Calpain-mediated cleavage of Atg5 switches autophagy to apoptosis. Nat Cell Biol 8, 1124–1132 (2006).

29. Lee, J.L. & Streuli, C.H. Integrins and epithelial cell polarity. J Cell Sci 127, 3217–3225 (2014).

30. Tsujimura, A. et al. Proximal location of mouse prostate epithelial stem cells: a model of prostatic homeostasis. J Cell Biol 157, 1257–1265 (2002).

31. Karthaus, W.R. et al. Identification of multipotent luminal progenitor cells in human prostate organoid cultures. Cell 159, 163–175 (2014).

32. Wang, X. et al. A luminal epithelial stem cell that is a cell of origin for prostate cancer. Nature 461, 495–500 (2009).

33. Liu, K. et al. Triple-cell lineage tracing by a dual reporter on a single allele. J Biol Chem 295, 690–700 (2020).

34. Guo, W. et al. JAK/STAT signaling maintains an intermediate cell population during prostate basal cell fate determination. Nat Genet 56, 2776–2789 (2024).

35. Bentsen, M. et al. ATAC-seq footprinting unravels kinetics of transcription factor binding during zygotic genome activation. Nat Commun 11, 4267 (2020).

36. Olive, M. et al. A dominant negative to activation protein-1 (AP1) that abolishes DNA binding and inhibits oncogenesis. J Biol Chem 272, 18586–18594 (1997).

37. Biddie, S.C. et al. Transcription factor AP1 potentiates chromatin accessibility and glucocorticoid receptor binding. Mol Cell 43, 145–155 (2011).

38. English, H.F., Santen, R.J. & Isaacs, J.T. Response of glandular versus basal rat ventral prostatic epithelial cells to androgen withdrawal and replacement. Prostate 11, 229–242 (1987).

39. Evans, G.S. & Chandler, J.A. Cell proliferation studies in the rat prostate: II. The effects of castration and androgen-induced regeneration upon basal and secretory cell proliferation. Prostate 11, 339–351 (1987).

40. Choi, N., Zhang, B., Zhang, L., Ittmann, M. & Xin, L. Adult murine prostate basal and luminal cells are self-sustained lineages that can both serve as targets for prostate cancer initiation. Cancer Cell 21, 253–265 (2012).

41. Kwon, O.J., Zhang, L., Ittmann, M.M. & Xin, L. Prostatic inflammation enhances basal-to-luminal differentiation and accelerates initiation of prostate cancer with a basal cell origin. Proc Natl Acad Sci U S A 111, E592–600 (2014).

42. Gleave, M.E. et al. Randomized comparative study of 3 versus 8-month neoadjuvant hormonal therapy before radical prostatectomy: biochemical and pathological effects. The Journal of urology 166, 500–506; discussion 506–507 (2001).

43. Taplin, M.E. et al. Effect of neoadjuvant abiraterone acetate (AA) plus leuprolide acetate (LHRHa) on PSA, pathological complete response (pCR), and near pCR in localized high-risk prostate cancer (LHRPC): results of a randomized phase II study. J Clin Oncol 30 (Suppl), Abstract 4521 (2012).

44. Dallos, M.C. et al. Androgen Deprivation Therapy Drives a Distinct Immune Phenotype in Localized Prostate Cancer. Clin Cancer Res 30, 5218–5230 (2024).

45. Tang, F. et al. Chromatin profiles classify castration-resistant prostate cancers suggesting therapeutic targets. Science 376, eabe1505 (2022).

46. Chan, J.M. et al. Lineage plasticity in prostate cancer depends on JAK/STAT inflammatory signaling. Science 377, 1180–1191 (2022).

47. Han, X. et al. A suite of new Dre recombinase drivers markedly expands the ability to perform intersectional genetic targeting. Cell Stem Cell 28, 1160–1176 e1167 (2021).

48. Li, F. et al. ERG orchestrates chromatin interactions to drive prostate cell fate reprogramming. J Clin Invest (2020).

49. Ni, Z., Chen, S., Brown, J. & Kendziorski, C. CB2 improves power of cell detection in droplet-based single-cell RNA sequencing data. Genome Biol 21, 137 (2020).

50. Wolf, F.A., Angerer, P. & Theis, F.J. SCANPY: large-scale single-cell gene expression data analysis. Genome Biol 19, 15 (2018).

51. Traag, V.A., Waltman, L. & van Eck, N.J. From Louvain to Leiden: guaranteeing well-connected communities. Sci Rep 9, 5233 (2019).

52. van Dijk, D. et al. Recovering Gene Interactions from Single-Cell Data Using Data Diffusion. Cell 174, 716–729 e727 (2018).

53. Dobin, A. et al. STAR: ultrafast universal RNA-seq aligner. Bioinformatics 29, 15–21 (2013).

54. Langmead, B. & Salzberg, S.L. Fast gapped-read alignment with Bowtie 2. Nat Methods 9, 357–359 (2012).

55. Zhang, Y. et al. Model-based analysis of ChIP-Seq (MACS). Genome Biol 9, R137 (2008).

56. Heinz, S. et al. Simple combinations of lineage-determining transcription factors prime cis-regulatory elements required for macrophage and B cell identities. Mol Cell 38, 576–589 (2010).

57. Liao, Y., Smyth, G.K. & Shi, W. featureCounts: an efficient general purpose program for assigning sequence reads to genomic features. Bioinformatics 30, 923–930 (2014).

58. Love, M.I., Huber, W. & Anders, S. Moderated estimation of fold change and dispersion for RNA-seq data with DESeq2. Genome Biol 15, 550 (2014).

